# JAK and TYK2 inhibitors differentially modulate interferon/TNF-driven inflammation, stemness and proliferation in the colonic epithelium of ulcerative colitis

**DOI:** 10.64898/2026.07.20.739113

**Authors:** Arun Sridhar, Gunnar Andreas Enerhaug Walaas, Siri Sæterstad, Johan Pierre Durand Myrmehl, Romana Cermakova, Marte Gisvold Myrseth, Leona Grundel, Marianne Doré Hansen, Marianne Otterstad, Marte Lie Høivik, Ann Elisabet Østvik, Ingunn Bakke, Torunn Bruland

## Abstract

**Background:** Janus kinase (JAK)-Signal Transducer and Activator of Transcription (STAT) pathway is a key regulator of inflammatory signaling in ulcerative colitis (UC). While most studies have focused on JAK/tyrosine kinase 2 (TYK2) inhibitors effects on immune cell-mediated responses, their direct epithelial impact remains less known. We investigated epithelial-specific transcriptional responses to JAK/TYK2 inhibitors using patient-derived intestinal epithelial organoids (IEOs) under UC-relevant conditions.

**Methods:** Colonic IEOs from UC patients were pretreated with various concentrations of tofacitinib, upadacitinib, filgotinib, brepocitinib, and deucravacitinib for 1 hour prior to stimulation with IFNβ, IFNγ, or IFNλ1 for Western blot analysis of STAT1/3 and TYK2 phosphorylation. For transcriptomic profiling, IEOs were pretreated with upadacitinib or deucravacitinib for 16 hours, followed by 8-hour stimulation with IFNγ, IFNλ1, TNF, or IFNγ + TNF. Bulk RNA sequencing assessed differential gene expression, and multiplex assays quantified chemokine secretion. Ki67 immunohistochemistry on colonic biopsies from healthy controls, and UC patients with and without JAK inhibitors-treatment were assessed for epithelial proliferation.

**Results:** IFNs induced distinct STAT1/3 and TYK2 activation, with IFNβ/γ eliciting stronger phosphorylation than IFNλ1. All JAK/TYK2 inhibitors regulated pSTAT1/3 and pTYK2, with upadacitinib most strongly inhibiting pSTAT1/3 and deucravacitinib selectively targeting pTYK2. Transcriptomic analysis revealed extensive cytokine-driven gene regulation, with IFNγ + TNF eliciting the strongest response. Enrichment analysis highlighted upregulation of IFN signaling, antigen presentation, and innate immune pathways, alongside downregulation of cell-cycle processes. Drug-response profiling showed minimal transcriptional changes with upadacitinib and deucravacitinib alone. Upadacitinib broadly modulated IFNs and IFNγ + TNF-regulated genes, attenuating JAK-STAT, NFκB, antiviral, and cell death pathways, while restoring genes linked to mucosal healing. Upadacitinib also reduced IFNs and IFNγ + TNF-driven chemokine genes and protein secretion. In contrast, deucravacitinib showed selective, potent inhibition of inflammatory genes under IFNλ1-stimulation. Both inhibitors minimally impacted TNF-driven pathways. Ki67 immunohistochemistry confirmed enhanced epithelial proliferation in JAK inhibitor-treated UC patients regardless of mucosal inflammation status.

**Conclusions:** Our findings provide novel evidence that JAK/TYK2 inhibitors influence epithelial transcriptional programs associated with inflammation and mucosal healing. Upadacitinib demonstrated broader modulation of cytokine-driven gene networks compared to TYK2-selective inhibition. These findings provide insight into epithelial-specific drug actions and support precision approaches for UC therapy.

## Background

Ulcerative colitis (UC), a major subtype of inflammatory bowel disease (IBD), is characterized by chronic, relapsing inflammation of the colonic mucosa, with multifactorial etiology (1). The pathogenesis of UC is likely driven by a complex interplay between genetic susceptibility, environmental factors and dysregulated microbiome-host interactions, with epithelial barrier dysfunction emerging as a central contributor to disease progression (2). Neutrophil infiltration and elevated secretion of pro-inflammatory cytokines including interferon-gamma (IFNγ), tumor necrosis factor (TNF), and interleukins (IL-5, IL-6, IL-13, IL-17, and IL-22), significantly contributes to the mucosal damage and impairs epithelial homeostasis (3, 4). Current therapeutic strategies include conventional immunosuppressants, targeted biologic agents, and small molecule therapies such as Janus kinase (JAK) inhibitors and Sphingosine 1-phosphate (S1P) modulators (5–8). These therapies are designed to inhibit inflammation rather than promoting mucosal healing and restore intestinal homeostasis (9). However, a substantial subset of patients remains refractory to these interventions. Despite accumulating evidence indicating epithelial dysfunction in both disease onset and persistence (10, 11), drugs specifically targeting epithelial regeneration and barrier restoration are still underdeveloped (12, 13).

JAK inhibitors have emerged as a promising therapeutic modality due to their capacity to simultaneously modulate multiple cytokines (14). The JAK-Signal Transducer and Activator of Transcription (STAT) pathway is an evolutionarily conserved signaling pathway and a crucial mediator in IBD pathogenesis (15). This pathway comprises four intracellular non-receptor tyrosine kinases (JAK1, JAK, JAK3 and tyrosine kinase 2 (TYK2)) and seven STAT family members (STAT1, STAT2, STAT3, STAT4, STAT5A, STAT5B, and STAT6) (16). Activation of the JAK-STAT pathway occurs through inflammatory cytokines including IFNs and the interleukin families IL-6, IL-9, IL-10, and IL-12/23 (17–20). Interferons have multifaceted role in the intestine, acting as a critical regulatory node for antiviral defense, maintenance of intestinal homeostasis and modulation of inflammatory responses in IBD pathophysiology (19, 21). Type II IFN (IFNγ) serves as an immune modulator important for resistance to many microbial infections, while type I (e.g., IFNα, IFNβ) and type III (IFNλ subtypes) share similar induction mechanisms, stimulate interferon-stimulated genes (ISGs), and are mostly recognized for their ability to restrict virus infections (22). However, type-I IFNs can activate nearly all nucleated cells, whereas type III IFN responses are limited to few cell types, including epithelial cells (21, 22). The JAK-STAT transduction cascade is initiated when each IFN type binds to its specific receptor complex, each of which is associated with distinct combination of JAK kinase. Interferon signaling in intestinal epithelial cells (IECs) has been reported to exert both protective and detrimental effects (21, 23, 24).

Currently, several JAK/TYK2 inhibitors are approved or in various stages of clinical development for IBD. Among these, tofacitinib (25) (oral pan-JAK inhibitor), upadacitinib (26) and filgotinib (27) (oral JAK1-specific inhibitors) have received regulatory approval for moderate-to-severe UC patients. Given the potentially reduced toxicity profile compared to JAK1-3 inhibitors, the more recently developed TYK2 inhibitors such as deucravacitinib (28) (TYK2 inhibitor) and brepocitinib (29) (JAK1/TYK2 inhibitor) are under investigation as potential therapeutic options. The different IFN types differ in their requirements of JAKs and STATs indicating that the pan-JAK and the JAK-specific inhibitors will have different effects on the cell depending on which ligand-receptor complex is activated (19, 30). This differential selectivity confers potential therapeutic advantages; JAK1-specific inhibition may effectively suppress IFNγ-mediated inflammatory cascades while TYK2 inhibition could preserve beneficial epithelial repair mechanisms mediated through alternative JAK pathways. Further, these small molecules may exhibit different affinity profiles and function by inhibiting specific JAKs, thereby selectively modulating specific inflammatory arms of epithelial JAK-STAT signaling while preserving essential homeostatic functions. The IECs play a critical role in IBD pathophysiology (31) and are reported to respond to diverse pro-inflammatory stimuli by inducing inflammatory genes in the JAK-STAT and NFκB pathways, antigen presentation genes (32, 33), cell death (34–38), antiviral protection system (34), barrier function disruption (39, 40), and regulation of mucosal healing (39, 41, 42). JAK inhibitors can modulate these inflammatory pathways mediated by cytokines in immune as well as non-immune cell types (7, 43). Furthermore, multiple IBD-associated genes encode or regulate components of this JAK pathway, highlighting its genetic significance in disease etiology (44–46). However, inhibition of IFN signaling may compromise innate antiviral defense mechanisms, as evidenced by increased infection rates observed with JAK inhibitors in therapies (47).

Despite the central roles of the intestinal epithelium in mucosal healing, the cellular and molecular mechanisms underlying its responses to JAK inhibitors remain incompletely understood. Patient-derived intestinal epithelial organoids (IEOs) closely mimic three-dimensional intestinal epithelial architecture and maintain the genetic profiles of the donor, facilitating investigation of patient-specific responses and drug mechanisms of action (48, 49). Therefore, this study aims to delineate the effects of first- and next-generation JAK/TYK2-specific inhibitors using patient-derived colonic IEOs treated with UC-relevant cytokines.

## Methods

Experimental overviews are presented in Figure 1A, Figure 2A and Supplementary Figure 1A. Characteristics of donors used for IEO generation and for *in vivo* immunohistochemistry staining are given in Tables 1, 2 and Supplementary File 1, respectively.

**Figure 1:**
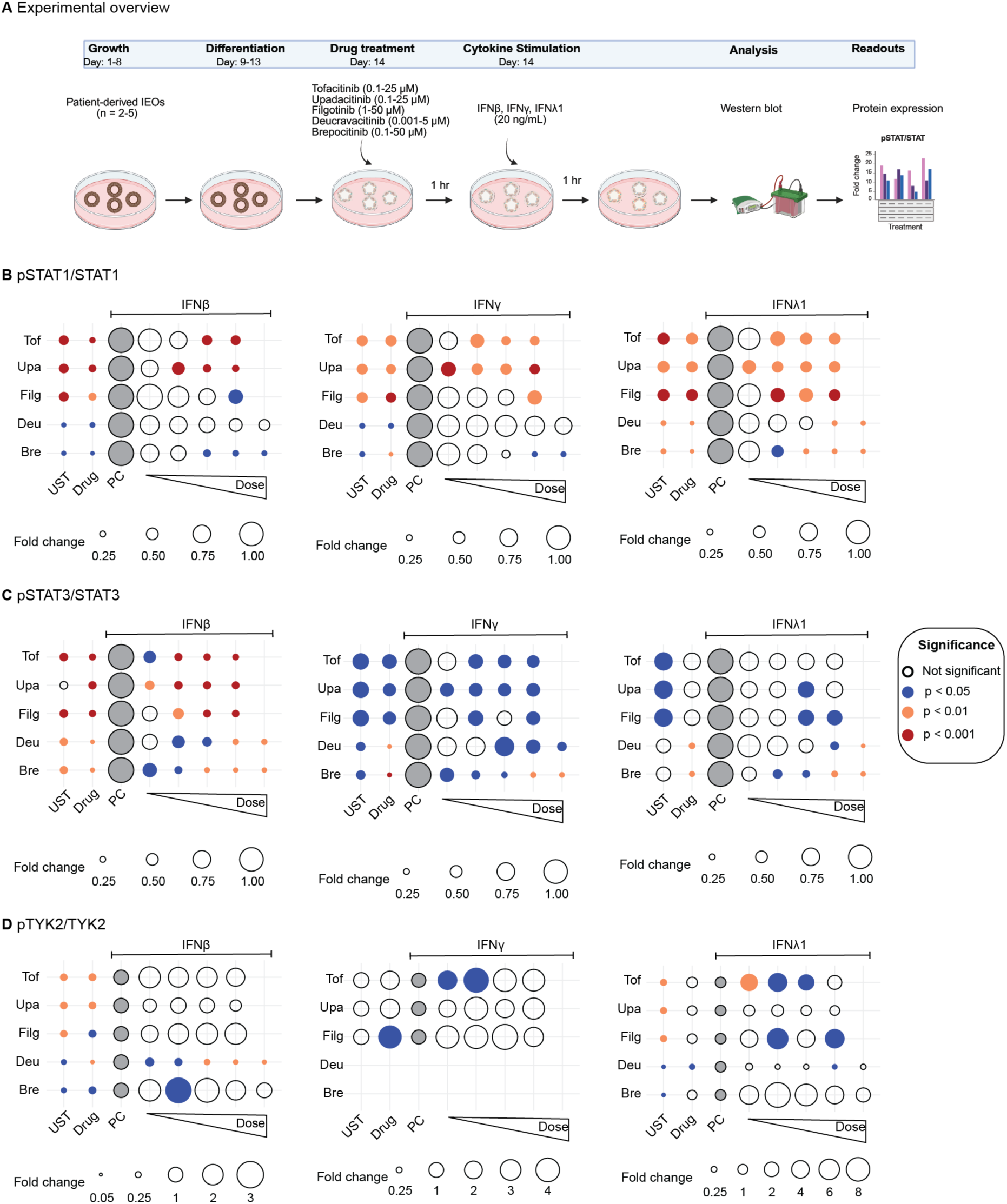
Experimental design and impact of JAK-TYK2 inhibitors on IFN-induced pSTAT1, pSTAT3, and pTYK2 in IEOs. A) Schematic illustration of the experimental workflow used for immunoblotting analysis. IEOs (*n* = 2-5) were pretreated for one hour with either DMSO (vehicle control) or tofacitinib (0.1, 1, 10, 25 µM), upadacitinib (0.1, 1, 10, 25 µM), filgotinib (1, 10, 25 and 50 µM), deucravacitinib (0.001, 0.025, 0.1, 1 and 5 µM), and brepocitinib (0.1, 1, 10, 25 and 50 µM). This was followed by stimulation with IFNβ, IFNγ, and IFNλ1 (20 ng/mL) for one hour, where applicable. B-D) Statistical effects of the JAK/TYK2 inhibitors on pSTAT1, pSTAT3 and pTYK2 are presented in bubble graphs where fold change is calculated by normalization to its respective interferon-stimulated control (PC) (grey circles), followed by normalizing to GAPDH. Statistical significance was assessed using repeated-measures one-way ANOVA followed with Holm-Sidak’s multiple post hoc test. Bubble size represents fold change, with each color showing significance level: No fill (no significance), blue fill (p < 0.05), orange fill (p < 0.01), and red fill (p < 0.001). Tof – tofacitinib, Upa - upadacitinib, Filg – filgotinib, Deu – deucravacitinib and Bre – brepocitinib. Representative blots and quantification of fold changes are shown in Supplementary Figures 2-4; full blots in Supplementary figures 12-15.

**Figure 2:**
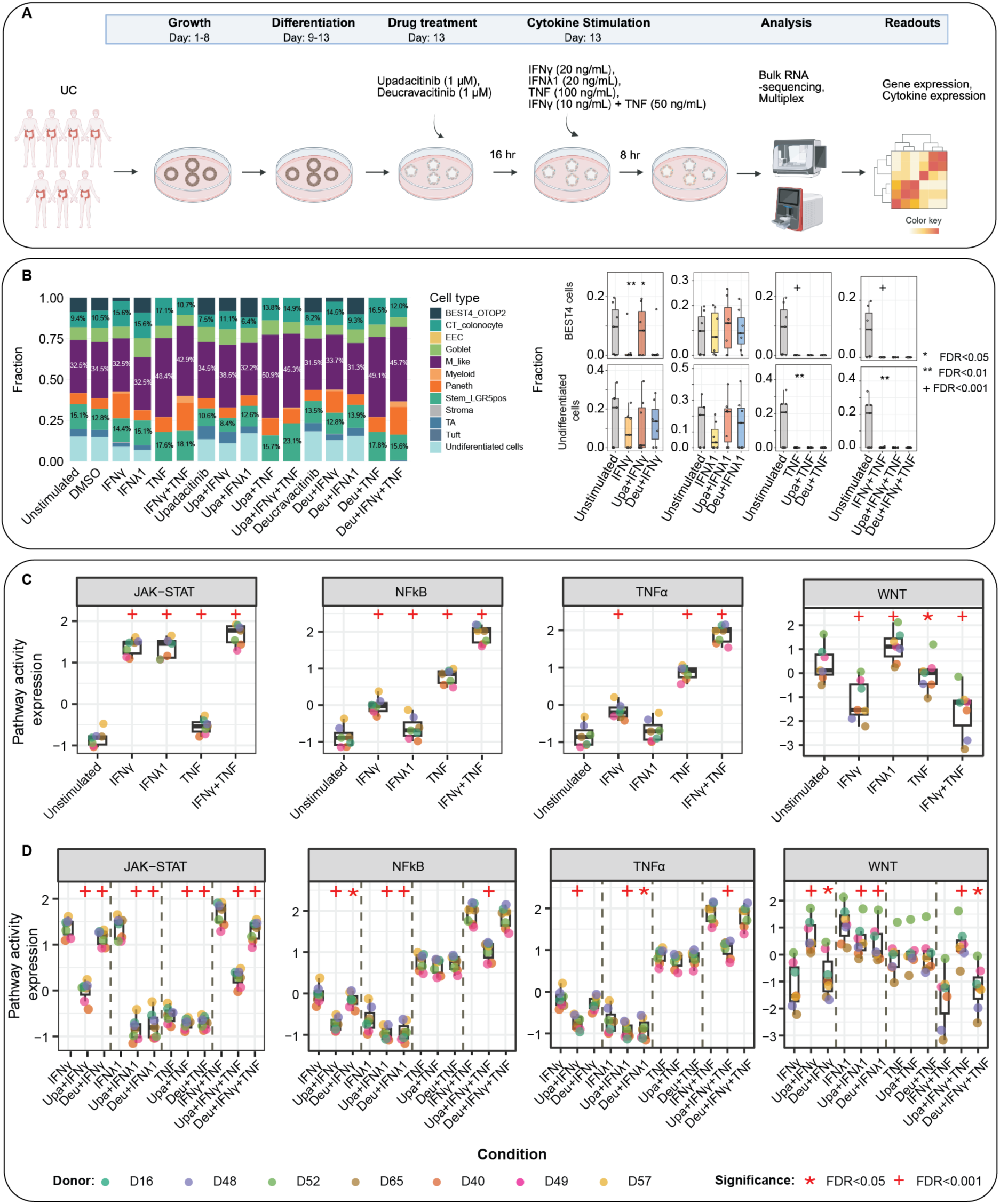
Upadacitinib inhibits JAK-STAT, NFκB, and TNF pathways and restores WNT signaling. A) Graphical representation of experimental design used for RNA sequencing and multiplex cytokine profiling. B) Estimated fractions of epithelial cell types in IEOs by CIBERSORTx deconvolution. Box plots show the fraction of BEST4+ and undifferentiated epithelial cells from 7 donors across the indicated conditions. Significance was assessed by linear mixed models (LMM) on centered log-ratio-transformed proportions with donor as a random intercept and P-values adjusted using the Benjamini-Hochberg method *FDR < 0.05, **FDR < 0.01, **^+^**FDR < 0.001 C,D) Box plots shows the pathway activity expression in IEOs (n = 7), inferred using PROGENy (65), from variance-stabilized transformed RNA-seq counts. for C) each stimulated condition compared to unstimulated control and D) each JAK/TYK2 inhibitor pretreatment followed by stimulation compared to their matched stimulation alone. Statistical significance was assessed by LMM with donors treated as a random effect. P-values were adjusted using Benjamini-Hochberg method for multiple comparisons testing. *FDR<0.05, **^+^**FDR<0.001. Upa – Upadacitinib; Deu - Deucravacitinib.

**Table 1:**
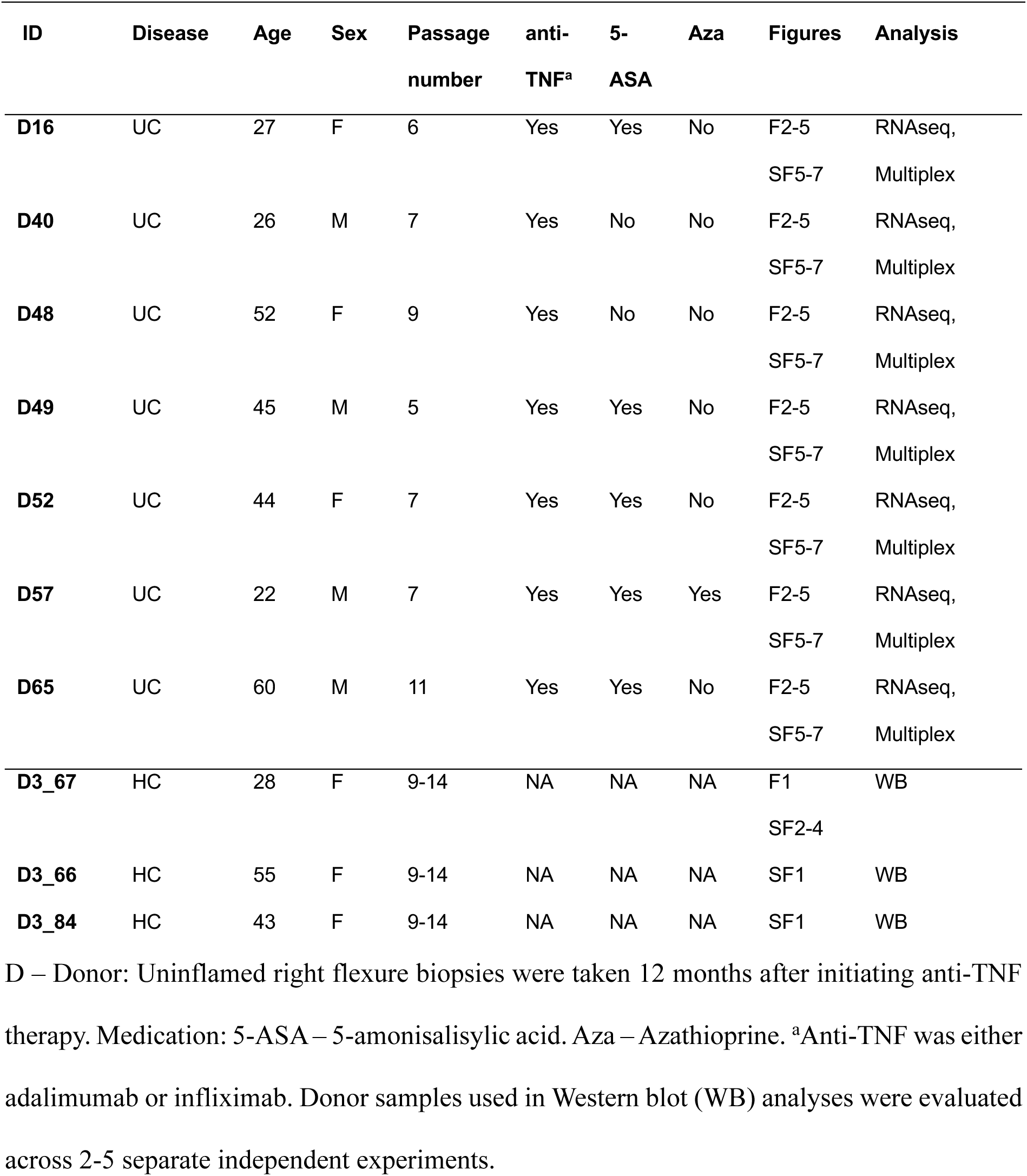
Patient characteristics (IEO generation)

| ID | Disease | Age | Sex | Passage<br>number | anti-TNF<sup>a</sup> | 5-ASA | Aza | Figures | Analysis |
| --- | --- | --- | --- | --- | --- | --- | --- | --- | --- |
| **D16** | UC | 27 | F | 6 | Yes | Yes | No | F2-5<br>SF5-7 | RNAseq,<br>Multiplex |
| **D40** | UC | 26 | M | 7 | Yes | No | No | F2-5<br>SF5-7 | RNAseq,<br>Multiplex |
| **D48** | UC | 52 | F | 9 | Yes | No | No | F2-5<br>SF5-7 | RNAseq,<br>Multiplex |
| **D49** | UC | 45 | M | 5 | Yes | Yes | No | F2-5<br>SF5-7 | RNAseq,<br>Multiplex |
| **D52** | UC | 44 | F | 7 | Yes | Yes | No | F2-5<br>SF5-7 | RNAseq,<br>Multiplex |
| **D57** | UC | 22 | M | 7 | Yes | Yes | Yes | F2-5<br>SF5-7 | RNAseq,<br>Multiplex |
| **D65** | UC | 60 | M | 11 | Yes | Yes | No | F2-5<br>SF5-7 | RNAseq,<br>Multiplex |
| **D3_67** | HC | 28 | F | 9-14 | NA | NA | NA | F1<br>SF2-4 | WB |
| **D3_66** | HC | 55 | F | 9-14 | NA | NA | NA | SF1 | WB |
| **D3_84** | HC | 43 | F | 9-14 | NA | NA | NA | SF1 | WB |

D – Donor: Uninflamed right flexure biopsies were taken 12 months after initiating anti-TNF therapy. Medication: 5-ASA – 5-aminosalicylic acid. Aza – Azathioprine. <sup>a</sup>Anti-TNF was either adalimumab or infliximab. Donor samples used in Western blot (WB) analyses were evaluated across 2-5 separate independent experiments.

**Table 2:** Patient characteristics (IHC validation cohort)

| Disease category | HC | UC without JAK inhibitor | UC with JAK inhibitors |
| --- | --- | --- | --- |
| **n** | 10 | 14 | 11 |
| **Sex** | 6M,4F | 11M, 3F | 9M, 2F |
| **Age**<br>(median+range) | 51 (25-69) | 43.5 (24-72) | 40 (23-69) |
| **Disease Duration** | NA | 13.5 (1-50) | 15 (1-33) |
| **Inflammatory status** | Uninflamed | 12 Uninflamed,<br>2 mild chronically inflamed | 6 Uninflamed,<br>4 mild chronically inflamed,<br>3 severe chronically inflamed |
| **Tissue location** | Hepatic flexure | Hepatic flexure | 11 sigmoid<sup>#</sup>, 2 ascending colon<sup>#</sup> |
| **JAK inhibitors** | NA | No | Filgotinib (Jyseleca) /<br>Tofacitinib (Xeljanz) |
| **Other medication** | NA | One patient using 5-ASA, 5<br>patients using local 5-ASA<br>rectum | One patient using 5-ASA |

F – Female; HC – healthy control; M – Male; NA – Not applicable; UC – Ulcerative colitis; Medication: 5-ASA – 5-aminosalicylic acid. <sup>#</sup>For two patients in the JAK inhibitor-treated group, biopsies were taken from both ascending and sigmoid colon with differing local inflammation status, and each biopsy was analyzed as a separate observation reflecting its local tissue state. All remaining patients contributed single biopsy.

### Culturing and experiments in HT-29 cells

HT-29 cells from ATCC (#HTB-38, RRID:CVCL_0320) were cultured at 37°C, 5% CO2 in RPMI medium (#R8758, Sigma-Aldrich), supplemented with10% Fetal Bovine Serum (FBS, #10270106, Thermo Fischer Scientific), 2mM L-Glutamine (#G7513, Sigma-Aldrich) and 0.05 mg/mL Gentamicin (#G1397, Sigma-Aldrich). For experiments, confluent cells were seeded in two 6-well plates (# 3518, Corning) and incubated 24 hours in FBS-depleted (5%) starvation medium to inhibit growth, before stimulation with either 10 ng/mL IFNβ (#300-02BC, PeproTech), IFNγ (#300-02, Peprotech) or IFNλ1 (#300-02L, PeproTech) for the indicated times (15-240 minutes). After the incubation, cells were harvested on ice by washing with cold Dulbecco’s Phosphate Buffered Saline (DPBS, #8537, Sigma-Aldrich) before detaching using cell scraper and 80µl lysis buffer/well, and stored at -20°C for measurement of protein concentration and immunoblot.

### Culturing and experiments in intestinal epithelial organoids (IEOs)

Patient inclusion criteria and clinical characteristics have been described previously (50) and are summarized in Table 1. IEOs were derived from uninflamed colonic (right flexure) pinch biopsies (4-5 biopsies per patient) and were established and maintained following a protocol adapted from Jung et al. (51) and Mahe et at. (52), with modification as described in Gopalakrishnan et al. (53). We expanded the cells at 5% oxygen, and conducted experiments at 2% oxygen since stem cells and differentiated epithelial cells are adapted to slightly different levels of physiological hypoxia (54, 55). For experimental procedures, IEOs were cultured, grown, and differentiated as previously detailed in Walaas et al., (50). Briefly, single cells dissociated from IEOs were resuspended in Matrigel from the same lot (#3018001, Corning, 7.7 mg/mL protein) at a concentration of 9000 cells/50μL and plated on prewarmed 24-well plates (2x25μL domes/well). IEOs were grown until day 8 and on day 9, culturing media was altered to differentiation media containing 5% WNT and DAPT (Notch signaling inhibitor)(53). This medium was refreshed on days 10 and 13. For investigating the activation pattern of pSTAT1/3 and pTYK2 in response to interferon-stimulation, IEOs were either treated with 20 ng/mL of IFNβ, IFNγ or IFNλ1 (#300-02BC, #300-02 or #300-02L, PeproTech) for 15 and 60 minutes. To assess the JAK inhibitors effect, IEOs were pretreated for one hour with 0.1, 1, 10, 25 μM tofacitinib (CP-690550, #S5001, Selleckchem), 0.1, 1, 10, 25 μM upadacitinib (#ABT-494, Selleckchem), 1, 10, 25 and 50 μM of filgotinib (#GLP0634, Selleckchem), 0.001, 0.025, 0.1, 1 and 5 μM deucravacitinib (#S8879, Selleckchem) and 0.1, 1, 10, 25 and 50 μM brepocitinib (#S8804, Selleckchem), followed by one hour of stimulation with 20 ng/mL of IFNβ, IFNγ or IFNλ1. For RNA sequencing of targeted JAK1/TYK2 inhibition of inflammatory responses, the IEOs were pretreated with 1 μM of either the JAK1-inhibitor upadacitinib or the TYK2-inhibitor deucravacitinib for 16 hours, followed by 8 hours simultaneous stimulation with cytokines to reproduce the inflammatory environment of active UC: TNF (100 or 10 ng/mL, #300-01A, PeproTech), IFNγ (20 or 2 ng/mL), IFNλ1 (20 ng/mL), and IFNγ + TNF (10 ng/mL + 50 ng/mL). For each experiments using JAK/TYK-inhibitor drugs, dimethyl sulfoxide (DMSO) served as vehicle control at the highest relevant concentration. Post-stimulation, conditioned media was harvested and stored for multiplex profiling. The cells were either collected for immunoblotting as described previously (53) and stored at -80°C or lysed for RNA analyses by washing the cells with prewarmed PBS and adding 250μL ice cold lysis buffer with beta-mercaptoethanol. The Matrigel domes were scraped, resuspended, and transferred to a 2mL Eppendorf tube. Lastly, the lysates were passed through sterile 20-gauge needles ten times to get a homogenous lysate before they were stored at -80°C. Each condition had four to six technical replicates that were combined depending on the experiment.

### Immunoblotting

Proteins were isolated from the IEOs by lysing for 2 hours on ice using lysis buffer containing 50 mM Tris-HCl pH 7.5, 150 mM NaCl, 5 mM EDTA (#15575-038, Invitrogen), 1% NP-40 (#492018, Sigma-Aldrich), 1 mM dithiothreitol, 1x Complete® EDTA-free protease inhibitor (#11873580001, Sigma-Aldrich), and 1x phosphatase inhibitor cocktail II (#P5726, Sigma-Aldrich) and III (#P0044, Sigma-Aldrich), respectively. Cell lysates were clarified by centrifugation at 13,000 × g for 20 minutes at 4°C, and protein concentration was measured using the Pierce BCA protein assay (#23225, Thermo Fisher Scientific). Protein lysates were then denatured in 1 × NuPage lithium dodecyl sulfate sample buffer (#NP0007, Invitrogen) supplemented with 50 mM dithiothreitol for 10 minutes at 70°C before they were separated on 4–12% NuPage Bis-Tris gels (#NP0321BOX, Invitrogen) and electroblotted onto nitrocellulose membranes (0.2 μm, #1704158, Bio-Rad) using the Trans-Blot Turbo Transfer System (Bio-Rad). Overnight dried membranes were blocked using Rockland Blocking Buffer for Fluorescent Western Blotting (#MB-070, Rockland) for 1 hour at room temperature before incubation with the indicated antibodies overnight at 4°C. The blots were incubated with Dylight conjugated secondary antibodies for detection: Goat anti-Rabbit DyLight^TM^ 800 (1:5000, #SA5-35571) and Goat anti-Mouse DyLight^TM^ 680 (1:5000, #35518) (both from Invitrogen). Images were obtained with LI-COR Odyssey Imager and analyzed using Image Studio Software (LI-COR Biosciences). Total protein levels were normalized to glyceraldehyde-3-phosphate dehydrogenase (GAPDH) expression and given as fold change relative to untreated DMSO/positive control (Supplementary File 2). The following antibodies (all from Cell Signaling Technology) and dilutions were used: pSTAT1 (D4A7) rabbit monoclonal antibody (1:1000, #7649), STAT1-total (9H2) mouse monoclonal antibody (1:1000, #9176), pSTAT3 (D3A7) rabbit monoclonal antibody (1:1000, #9145), STAT3-total (D3Z2G) rabbit monoclonal antibody (1:1000, #12640), pTYK2 (D7T8A) rabbit monoclonal antibody (1:1000, #68790), TYK2 (D4I5T) rabbit monoclonal antibody (1:1000, #14193) and GAPDH (D16H11) rabbit monoclonal antibody (1:5000, #5174).

### RNA isolation and sequencing

RNA was extracted from IEOs and processed for sequencing as previously described in Walaas et al., (50). Briefly, total RNA was isolated using the RNeasy mini kit (#74106, Qiagen) and concentration and quality were evaluated with NanoDrop Spectrophotometer (Thermo Scientific). RNA sequencing libraries were prepared using the Illumina Stranded mRNA Prep (Ligation) Kit, following the manufacturer’s protocol. Sequencing was performed on the NovaSeq 6000 platform. Data processing including quality control with FastQC (v0.11.9), and fastp (v0.20.1), alignment using the STAR aligner (v2.7.9a), and transcript quantification was carried out using quasi-alignment with Salmon (v1.7.0). Differential expression analysis was conducted using the DESeq2 Bioconductor package (56).

### Cell deconvolution

Colonic single-cell RNA sequencing data from Thomas et al. (57) (815,900 cells; GSE282122), restricted to the epithelial compartment, was used as the reference dataset. Cell state-specific reference profiles were generated from pseudobulk expression profiles normalized to counts per million (CPM) and used to construct a custom signature matrix in CIBERSORTx (58). Deconvolution of IEO bulk RNA-seq samples from 7 donors was performed using the CIBERSORTx fractions module with B-mode batch correction and quantile normalization disabled, consistent with RNA-seq input data (58). Estimated fractions were analyzed as compositional data. Zero values were replaced using a pseudocount equal to half of the smallest non-zero fraction observed across the dataset, followed by centered log-ratio (CLR) transformation. For each inferred epithelial state, differences across treatment conditions (unstimulated, IFNγ, IFNλ1, TNF, and IFNγ + TNF stimulation, with or without co-treatment using upadacitinib or deucravacitinib) were modeled using linear mixed model on CLR-transformed values. The model included treatment condition as a fixed effect and donor as a random intercept to account for inter-individual variability [CLR ∼ Condition + (1|Donor)] (lme4) (59). P-values were adjusted across inferred epithelial states within each contrast using the Benjamini-Hochberg procedure, with significance defined as FDR < 0.05. Raw estimated fractions are displayed in figures, while statistical significance is derived from CLR-transformed mixed models.

### Multiplex Chemokine Profiling

The Bio-Plex Pro Human Chemokine Panel, 40-plex (#171AK99MR2, Bio-Rad) and the Bio-Plex 200 Systems were used to analyze undiluted IEO conditioned medium samples according to the manufacturer’s instructions.

### Immunohistochemistry

For *in vivo* assessment of epithelial proliferation, biopsies from ascending and sigmoid colon of UC patients receiving JAK inhibitory therapy (*n* = 11 (13 biopsies); *n* = 8 filgotinib (Jyseleca)/*n* = 3 tofacitinib (Xeljanz)) were obtained from the OUS IBD-biobank and research registry, Department of Gastroenterology, Oslo University Hospital, Norway. Two comparing groups of healthy controls (HC, *n* = 10) and UC patients not receiving JAK inhibitor therapy (*n* = 14) were obtained from a biobank at St. Olav’s Hospital/NTNU, Trondheim, Norway. The groups were matched for age, gender, disease duration and use of other medication (Supplementary File 1). Inflammation status was assessed by a pathologist.

Ki67 immunohistochemistry was performed on 4 µm formalin-fixed paraffin-embedded sections. After deparaffinization and rehydration, heat-induced epitope retrieval was performed with EnVision FLEX Target Retrieval Solution Low pH (Dako K8005) at 97 °C for 20 min in a PT Link module (Dako). Immunostaining was performed on a Dako Autostainer Plus using the Dako REAL EnVision Detection System Peroxidase/DAB+ rabbit/mouse kit (K8005). The primary anti-human Ki67 antibody (clone MIB-1, Dako, M72) was diluted 1:100 in Dako REAL Antibody Diluent (S2022) and incubated for 40 min at room temperature. Sections were counterstained with hematoxylin, dehydrated, and mounted with TissueMount (Sakura). Slides were scanned at 40× magnification on an Olympus SlideView VS200.

Tissue compartments (epithelium vs non-epithelium) were segmented using a DeepLabV3+ semantic segmentation model with a ResNet-18 encoder. Nuclei within epithelial regions were segmented using the StarDist 2D extension with a pretrained model (60). Ki67 positivity was then assigned to each segmented cell using a pretrained QuPath object classifier based on a DAB optical-density threshold in the nuclear compartment. Statistical analyses were performed in R software. Group differences were assessed by unpaired Wilcoxon rank-sum tests (Mann-Whitney U) with Benjamini-Hochberg correction across prespecified comparisons. Adjusted P-value (FDR) < 0.05 were considered significant.

### Bioinformatics and statistical analyses

Statistical analyses were performed using R software (V4.4) (RRID:SCR_001905) (61) or GraphPad Prism 9.0 (RRID: SCR_002798). Differential gene expression analysis resulted in multiple gene lists that were analyzed for intersecting patterns using the UpsetR and Euler packages (62, 63). Additional data visualization was performed using ggplot2 (64). Pathway activity inference was performed using PROGENy (65), based on variance-stabilized transformed expression counts data. To evaluate the statistical significance of pathway activity changes across conditions, we applied a linear mixed model (LMM) in R software using the Lmer4 (59) and LmerTest (66). Model formula: Expression ∼ Condition + (1|Donor), where ‘Condition’ was treated as fixed effect and ‘Donor’ as a random effect to account for inter-individual variability. Over-representation enrichment analyses was conducted using Metacore (Clarivate) and supported by DAVID (67, 68) and gprofiler2 (69). Input gene sets were defined using differential expression thresholds (log2 Foldchange > 0.03 and adjusted p-value < 0.05). Drugs effects on cytokine secretion were statistically evaluated with LMM: Log (Value) ∼ Condition + (1|Donor), where ‘Value’ represents the chemokine concentration, ‘Condition’ was the fixed effect, and ‘(1|Donor)’ accounts for random effects due to individual donor variation. P-value were adjusted using Benjamin-Hochberg method for multiple comparisons testing. Only chemokines with concentrations between 10 pg/ml to upper detection limit were analyzed. All the datasets with adjusted p < 0.05 were considered significant.

## Results

### Type I, II, and III interferons induce phosphorylation of STAT1, STAT3 and TYK2 in colonic epithelial cells

To systematically evaluate how different interferon types activate STAT1, STAT3 and TYK2 (70) in colonic IECs, we examined their phosphorylation patterns in response to 10 ng/mL IFNβ (type I), IFNγ (type II), and IFNλ1 (type III) by western blot. Initial experiments in HT-29 cells were conducted to determine the optimal timepoint for detection (experimental overview in Supplementary Figure 1A, upper panel). STAT1 and STAT3 phosphorylation were monitored over 15 to 240 minutes, while TYK2 phosphorylation was assessed at 15 and 120 minutes. In HT-29 cells, both IFNγ and IFNβ induced stronger STAT1 phosphorylation at 30 mins compared to IFNλ1 (Supplementary Figure 1B - left panel). STAT3 phosphorylation was most pronounced with IFNβ stimulation at 15 minutes followed by IFNγ at 60 minutes, while IFNλ1 induced the weakest response (Supplementary Figure 1B - center panel). TYK2 phosphorylation was strongest in response to IFNβ at 15 minutes, whereas IFNλ1 and IFNγ induced comparatively low phosphorylation (Supplementary Figure 1B - right panel).

To validate these findings in a human IEO model, we examined the same responses by western blot in IEOs (n=2) stimulated with 20 ng/mL IFNβ, IFNγ, and IFNλ1 at 15 and 60 minutes (experimental overview in Supplementary Figure 1A, lower panel). In IEOs, IFNβ induced the strongest phosphorylation, followed by IFNγ and IFNλ1 at 60 minutes for all three proteins STAT1, STAT3 and TYK2 (Supplementary Figure 1C – left, center and right panel, respectively). Together, these results demonstrate that IFNβ consistently induces the strongest activation of STAT1, STAT3 and TYK2 in both HT-29 cells and IEOs, while IFNγ and IFNλ1 showed distinct and generally weaker activation profiles.

### JAK/TYK2-specific inhibitors dose dependently regulate IFN-induced STAT1/3 and TYK2 phosphorylation in UC patient-derived IEOs

Given the robust, interferon-mediated activation of STATs and TYK2 phosphorylation (Supplementary Figure 1), we investigated the potency and specificity of the JAK-specific inhibitors tofacitinib, upadacitinib, filgotinib, the dual JAK/TYK2-specific brepocitinib and TYK2-specific deucravacitinib in UC patient-derived IEOs. IEOs were pretreated for one hour with varying concentrations of tofacitinib and upadacitinib (0.1-25 µM), filgotinib (1-50 µM), brepocitinib (0.1-50 µM) and deucravacitinib (0.001-5 µM) followed by one hour stimulation with IFNβ (Supplementary Figure 2), IFNγ (Supplementary Figure 3) or IFNλ1 (Supplementary Figure 4) (Experimental design in Figure 1A). To assess the potential direct effects of the drugs, highest concentration of each inhibitor alone was included alongside unstimulated controls, revealing no induction of STAT1, STAT3 or TYK2 phosphorylation (Supplementary Figure 2-4).

Upadacitinib demonstrated the highest potency among the tested inhibitors with significant inhibition on IFN-induced pSTAT1 at concentrations as low as 0.1 µM for IFNγ and IFNλ1, and 1 µM for IFNβ stimulation (Figures 1B). Tofacitinib required 1 µM to inhibit IFNγ and IFNλ1-induced pSTAT1 and 10 µM for IFNβ. Filgotinib consistently showed lowest potency, requiring 10 µM for IFNλ1 and up to 50 µM for IFNβ and IFNγ. Brepocitinib inhibited IFNλ1-induced pSTAT1 at 1 µM, IFNβ at 10 µM, and IFNγ at 25 µM. Deucravacitinib selectively inhibited IFNλ1-induced pSTAT1 at 1 µM, while showing a decreasing trend with IFNβ and IFNγ, suggesting a preferential impact on IFNλ1-mediated STAT1 activation. (Figures 1B).

Compared to pSTAT1 inhibition, more potent inhibition of IFN-induced pSTAT3 was observed substantially lower concentrations (Figures 1C). Upadacitinib significantly inhibited IFNβ and IFNγ-induced pSTAT3 levels at 0.1 µM, and at 10 µM for IFNλ1 (Figures 1C). Tofacitinib significantly inhibited IFNβ and IFNγ-induced pSTAT3 at 0.1 µM and 1 µM respectively but only showed a decreasing trend in IFNλ1-induced pSTAT3. Higher concentrations of filgotinib were required to significantly reduce IFN-induced pSTAT3; 10 µM for IFNβ and IFNγ and 25 µM for IFNλ1 (Figures 1C). Brepocitinib showed significant inhibition of pSTAT3 at 0.1 µM for IFNβ and IFNγ, and at 1 µM for IFNλ1 (Figure 1C). Notably, deucravacitinib exhibited potent and selective inhibition of pSTAT3 at concentrations as low as 0.025 µM for IFNβ, 0.1 µM for IFNγ and 1 µM for IFNλ1.

Regarding inhibition of TYK2 phosphorylation, the inhibitors displayed distinct inhibitory patterns (Figure 1D). Upon IFNβ stimulation, all JAK-specific inhibitors (tofacitinib, upadacitinib, and filgotinib) trended towards enhancing TYK2 phosphorylation though without reaching statistical significance (Figure 1D). Pronounced effects were observed with IFNγ, where tofacitinib significantly enhanced pTYK2 at 0.1 µM, despite IFNγ stimulation not directly inducing TYK2 receptors (Figure 1D). Similarly, with IFNλ1 stimulation, tofacitinib and filgotinib significantly enhanced pTYK2 at 0.1 µM and 10 µM, respectively (Figure 1D). The dual JAK/TYK2 inhibitor brepocitinib also increased pTYK2 with IFNβ at 1 µM and showed a trend towards activation with IFNλ1. In contrast, the TYK2-specific inhibitor deucravacitinib consistently reduced pTYK2 across all interferon stimulation, with significant inhibition at 0.001 µM for IFNβ and 0.1 µM for IFNλ1. Interestingly, upadacitinib neither induced nor inhibited IFNλ1-induced pTYK2 (Figures 1D).

Across all interferon stimulations, the inhibitors followed a consistent potency ranking of upadacitinib > deucravacitinib > brepocitinib > tofacitinib > filgotinib. Upadacitinib demonstrated the strongest inhibitory effect on both pSTAT1 and pSTAT3, while deucravacitinib uniquely suppressed pTYK2 (Figures 1B-D).

### Upadacitinib and deucravacitinib pretreatments exert strong transcriptomic changes in IFNs/TNF-stimulated IEOs, affecting a range of cellular processes

To further elucidate the mechanistic impact of targeted JAK1/TYK2 inhibition on epithelial inflammatory responses, we conducted comprehensive transcriptome profiling of IEOs pretreated with the JAK1-specific upadacitinib or TYK2-specific inhibitor deucravacitinib for 16 hours followed by stimulation with IFNγ, IFNλ1, TNF or IFNγ + TNF for 8 hours. This experimental design (Figure 2A) intended to mimic maintenance therapy condition (8), and capture drug effects on both early transcriptional responses and downstream gene regulation (71). IFNγ and IFNλ1 were selected to investigate their potential contrasting role in IBD inflammation: IFNγ is well-documented to have detrimental effects on IECs, especially in combination with TNF (7, 72), whereas IFNλ1, though less studied, has been reported to have protective function (23).

As recently reported (50), IFNs/TNF stimulation induced distinct transcriptional responses in IEOs, with IFNγ, IFNλ1 and TNF inducing 6,357, 1,069 and 4,070 differentially expressed genes (DEGs), respectively. The combination of IFNγ + TNF resulted in most pronounced response with 9,067 DEGs compared to unstimulated controls (Supplementary Figures 5, 6; Table 3). To quantify and compare cell type fractions in response to IFNs/TNF stimulation and after JAK/TYK2 inhibitor pretreatment, we performed deconvolution analysis using CIBERSORTx with a signature matrix built from colonic scRNA-seq data from Thomas et al. (57). Our signature matrix comprised of selected cell type signatures that included differentiated and undifferentiated epithelial subsets (colonocytes, transit amplifying cells, BEST4+/OTOP2+, goblet, paneth, LGR5+ stem, tuft, enteroendocrine cell, M-like, and undifferentiated cells) as well as myeloid and stromal compartments (Figure 2B). Myeloid and stromal signatures were significantly increased upon IFNγ/TNF stimulation, with the response appearing largely IFNγ-driven (Supplementary Figure 7, 8). Because our organoid system lacks immune and stromal cells, these increases most likely reflect inflammation-induced transcriptional signature of epithelial cells with mimics myeloid and stromal like states, rather than genuine shifts in cellular composition. Our results also show that upadacitinib pretreatment, which blocks IFNγ, restored the stromal signature under both IFNγ and IFNγ + TNF conditions and restored the myeloid fraction under IFNγ stimulation. whereas deucravacitinib did not significantly restore either signature. BEST4+ and undifferentiated cell signatures were significantly decreased upon IFNγ/TNF stimulation, and upadacitinib pretreatment restored only the BEST4+ signature (Figure 2B), while deucravacitinib failed to restore either. Overall, because inflammatory stimulation alters epithelial transcriptional state, these inferred fractions likely reflect both genuine shifts in cellular abundance and changes in epithelial transcriptional programs.

**Table 3:** List of upregulated and downregulated differential expressed genes (DEGs) after using a cutoff (|log2 foldchange| > 0.3 and p-adjusted < 0.05)

| Contrast | Upregulated DEGs | Downregulated DEGs | Total DEGs |
| --- | --- | --- | --- |
| Upa vs. DMSO | 1 | 11 | 12 |
| DEU vs. DMSO | 0 | 4 | 4 |
| Deu vs. Upa | 0 | 0 | 0 |
| IFNγ vs. Unstim | 2906 | 3451 | 6357 |
| Upa + IFNγ vs. IFNγ | 2917 | 2472 | 5389 |
| Deu + IFNγ vs. IFNγ | 7 | 56 | 63 |
| IFNλ1 vs. Unstim | 814 | 255 | 1069 |
| Upa + IFNλ1 vs. IFNλ1 | 374 | 995 | 1369 |
| Deu + IFNλ1 vs. IFNλ1 | 257 | 874 | 1131 |
| TNF vs. Unstim | 1801 | 2269 | 4070 |
| Upa + TNF vs. TNF | 1 | 8 | 9 |
| Deu + TNF vs. TNF | 0 | 7 | 7 |
| IFNγ + TNF vs. Unstim | 4237 | 4830 | 9067 |
| Upa + IFNγ + TNF vs. IFNγ + TNF | 3586 | 3402 | 6988 |
| Deu + IFNγ + TNF vs. IFNγ + TNF | 86 | 306 | 392 |
| IFNγ vs. IFNλ1 | 2483 | 3061 | 5544 |

Intestinal epithelial organoids (IEOs) from 7 donors were pretreated with the JAK1-specific upadacitinib (Upa) or TYK2-specific inhibitor deucravacitinib (Deu) for 16 hours prior to stimulation with IFNγ , IFNλ1, TNF or IFNγ + TNF for 8 hours. DEG analysis resulted in multiple gene lists that were analyzed further. Unstim – unstimulated.

To resolve the signaling programs driving these transcriptional shifts, we next assessed pathway activity using PROGENy (65), which revealed significant activation of JAK-STAT and NFkB pathways across all IFNs/TNF stimulation conditions (Figure 2C). Notably, IFNγ/TNF stimulation inhibited WNT signaling, while IFNλ1 activated it (Figure 2C). Additional pathways such as MAPK, EGFR and PI3K signaling were also significantly activated by all stimulations (Supplementary Figure 9). IFNγ, TNF and IFNγ + TNF significantly activated TNF (Figure 2C) and p53 signaling (Supplementary Figure 9), while TNF alone activated estrogen and hypoxia signaling. The TGFβ pathway was driven by IFNγ stimulation (Supplementary Figure 9). Over-representation analysis also revealed that processes such as interferon signaling, innate immune responses, antigen and phagosomes presentation, and G1-S phase interleukin regulation were consistently upregulated across IFNs/TNF-stimulated conditions (Supplementary Figures 5, 6). Processes related to cell cycle core, S phase, G2-M phase showed significant downregulation in IFNγ, TNF or IFNγ + TNF-stimulated conditions, while process related to DNA damages was primarily driven by TNF stimulation (Supplementary Figures 5, 6).

The transcriptional analysis further revealed that upadacitinib and deucravacitinib alone induced minimal transcriptional changes (12 and 4 DEGs respectively) (Table 3), while combining drugs with IFNs/TNF stimulation showed receptor specific anti-inflammatory effects. Upadacitinib demonstrated broad regulation of IFNγ-induced changes (5,389 DEGs) and deucravacitinib showed limited effect (63 DEGs). In contrast, both inhibitors showed comparable regulation of IFNλ1-induced transcriptional changes (1,131 vs 1,369 DEGs, respectively). Furthermore, both drugs had minimal effect on TNF-induced gene expression (7 vs 9 DEGs, respectively). Like IFNγ, under combined IFNγ + TNF stimulation, upadacitinib exhibited more extensive regulation with 6,988 DEGs compared to deucravacitinib with 392 DEGs (Supplementary Figures 5, 6; Table 3).

PROGENy analysis of pathway activities (65) following inhibitors pretreatment confirmed significant inhibition of the JAK-STAT pathway by both drugs across all IFNs/TNF-stimulated conditions (Figure 2D). Upadacitinib also inhibited NFkB and TNF pathways in response to IFNγ, IFNλ1 and IFNγ + TNF (Figure 2D), while deucravacitinib’s effects on these pathways were specific to IFNλ1 stimulation. Upadacitinib restored WNT signaling suppressed by IFNγ/TNF stimulation, but inhibited WNT activation induced by IFNλ1 stimulation. Neither drug affected TNF-induced pathways beyond JAK-STAT pathway. Effects on other pathways were generally non-significant and lacked specificity (Supplementary Figure 9). Over-representation enrichment analysis revealed that enriched processes such as interferon signaling, innate immune responses, antigen and phagosomes presentation, G1-S phase interleukin regulation were consistently downregulated by both upadacitinib or deucravacitinib pretreatment across IFNγ, IFNλ1, or IFNγ + TNF conditions (Supplementary Figures 5, 6). Processes downregulated during IFN/TNF stimulation such as cell cycle core, S phase, G2-M phase showed significant upregulation only with upadacitinib pretreatment while deucravacitinib pretreatment had minimal significant effect. Furthermore, neither of the drugs regulate any TNF induced DEGs. These data indicate that JAK1 inhibition through upadacitinib appears to be more effective at regulating IFNγ-mediated transcriptional changes. While TYK2 inhibition through deucravacitinib effectively inhibited inflammation-associated processes upregulated by IFNγ, IFNλ1, or IFNγ + TNF, it did not regulate cell cycle-related process networks and showed greater specificity for IFNλ1-mediated signaling.

### Upadacitinib and deucravacitinib differentially inhibit IFNs/TNF-induced inflammatory signaling pathways and chemokine release

Our data demonstrated that IFNγ, IFNλ1, TNF and IFNγ + TNF regulate an overlapping yet distinct pattern of gene expression in UC patient-derived IEOs. To further characterize the impact of JAK1/TYK2 inhibition, we conducted targeted analysis of inflammatory mediators within interferon pathways and NFκB signaling networks. This focused approach was based on our results from PROGENy analysis of pathway activities (65) and over-representation enrichment analysis and further guided by previously documented involvement of JAK-STAT (19, 73–75) and NFκB signaling (76, 77) in inflammatory contexts. Expression patterns in IEOs pretreated either with upadacitinib or deucravacitinib for 16 hours followed by 8 hours IFNs/TNF stimulation demonstrated robust differential expression across the JAK-STAT (Figure 3A) and NFκB signaling axes (Figure 3B). Notably, IFNγ + TNF combination elicited the most pronounced transcriptional changes followed by IFNγ and IFNλ1 (Figure 3). This ranked response pattern was evident throughout the receptor-mediated signaling networks.

**Figure 3:**
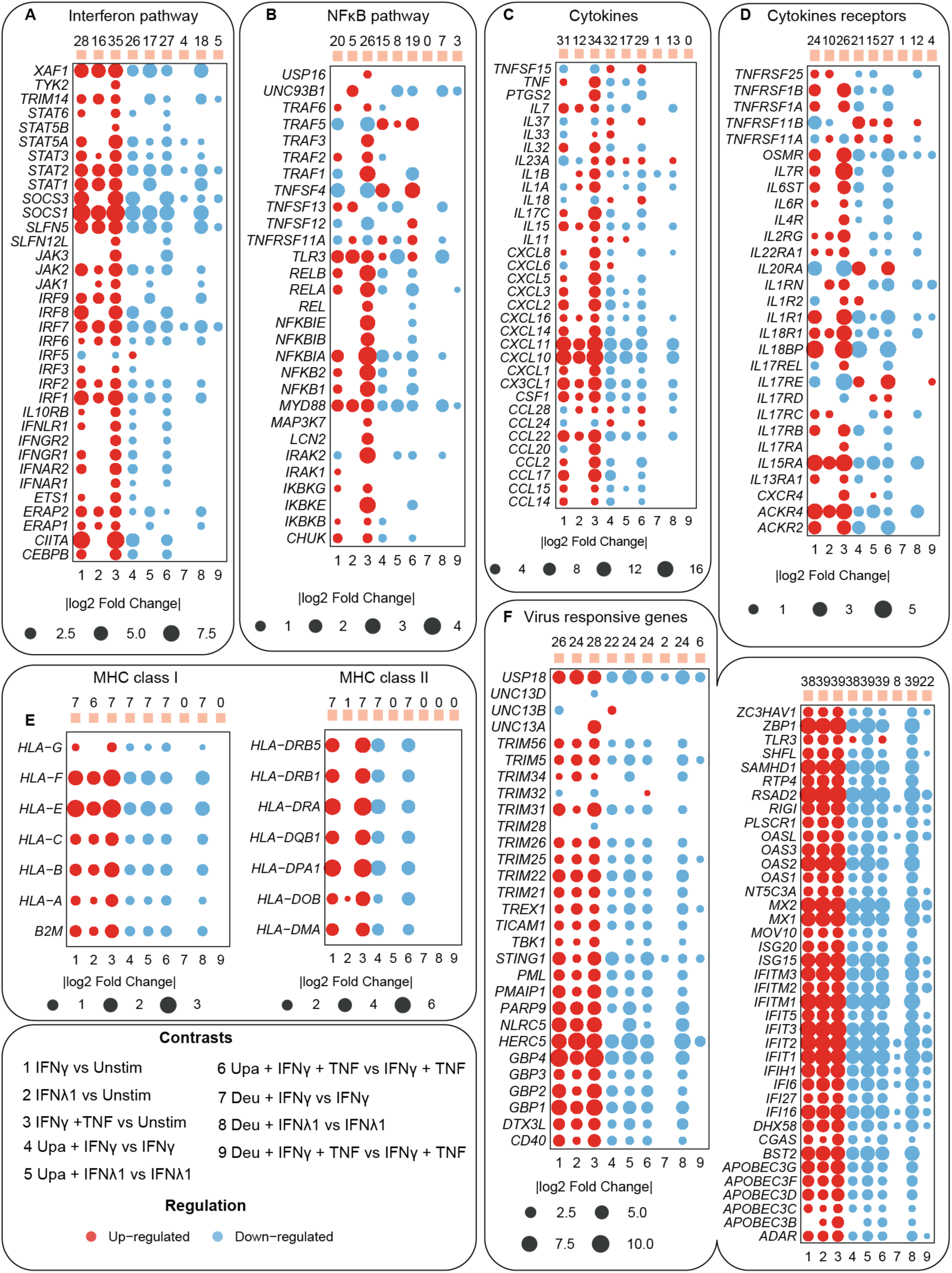
Upadacitinib and deucravacitinib decrease inflammation-associated genes expression in IFNγ or IFNγ + TNF-stimulated IEOs. Bubble plots showing log2 fold changes of selected genes in A) Interferon signaling, B) NFκB signaling, C) Cytokines, D) Cytokine receptors, E) MHC class I and II, and F) Viral responsive genes in IEOs (*n* = 7) stimulated with IFNγ, IFNλ1 and IFNγ + TNF (1–3) alone or together with pretreatment of upadacitinib (Upa) (4–6) or deucravacitinib (Deu) (7–9). Red and blue circles indicate up- and down-regulation, respectively, with circle size proportional to fold change. Only genes with p-value p < 0.05 are presented.

By examining genes involved in these inflammatory cascades, we observed stimulus-specific expression of cytokine receptors, signaling components and downstream effector genes. IFNγ + TNF combination uniquely upregulated both type I (*IFNAR1*, *IFNAR2*) and type II (*IFNGR1*, *IFNGR2*) interferon receptors, while simultaneously inducing TNF receptors (*TNFRSF1A*, *TNFRSF1B*) and multiple interleukin receptors (*IL1A*, *IL2RG*, *IL6R*, *IL6ST*, *IL7R*, *IL10RB*, *IL13RA1, IL15RA, IL17RB, IL18R1, IL18BP, IL22RA1*) (Figure 3A, D). This comprehensive receptor upregulation may establish a foundation for amplified signal transduction. In contrast, IFNγ alone selectively induced *IFNAR2, IFNGR1*, and a similar though less robust pattern of cytokine receptors, while IFNλ1 stimulation activated its specific receptors (*IL10RB, IFNLR1*) alongside a limited subset of interleukin receptors (*IL1RN, IL2RG, IL15RA, IL17RC, IL18R1, IL22RA1*) (Figure 3A, D).

IFNγ + TNF upregulated all JAKs (*JAK1, JAK2, JAK3* and *TYK2*) and multiple STAT transaction factors (*STAT1, 2, 3, 5A, 5B* and *6*). IFNγ selectively induced *JAK2* and a subset of STAT genes (*STAT1, 2, 3, 5A,* and *6*), while IFNλ1 activated *JAK1*, *JAK2*, and *STAT1, 2,* and *3*. We also observed stimulus-specific engagement of interferon regulatory factors (IRFs) with IFNγ and IFNγ + TNF combination upregulated *IRF1, IRF2, IRF3, IRF6, IRF7, IRF8, IRF9*, while IFNλ1 induced *IRF1, IRF2, IRF6, IRF7, IRF9*, and *CEBPB*. Both IFNγ and IFNγ + TNF combination downregulated *IRF5* (Figure 3A).

The JAK-STAT signaling networks were functionally integrated with NFκB pathway activation, particularly in IFNγ + TNF and IFNγ-stimulated conditions. These stimuli induced genes encoding the inhibitor of kappa B kinase (IκK) complex components (*CHUK* (IκK1), *IKBKB* (IκK2) and *IKBKG* (NEMO)) as well as interleukin-1-receptor associated kinases (*IRAK1, IRAK2*), and downstream NFκB components (*TLR3, NFΚB1, NFΚB2, MYD88, RELA, RELB, CEBPB, TRAF2*, and *TNFSF13*). Notably, *LCN2* induction was exclusive to IFNγ + TNF while IFNλ1 activated only *TLR3, MYD88*, and *TNFSF13*, reflecting a more limited activation of NFκB signaling (Figure 3B).

We also observed extensive activation of JAK-STAT and NFκB inducible chemokines and cytokine genes expression (Figure 3C). IFNγ + TNF stimulation induced the most robust induction of multiple cytokines families including CXC (*CXCL1, CXCL2, CXCL3, CXCL5, CXCL6, CXCL8, CXCL10, CXCL11, CXCL14, CXCL16*), CC (*CCL2, CCL15, CCL17, CCL20, CCL22*), CX3CL1, and ILs (*IL1A, IL1B, IL11, IL15, IL17C, IL23A, IL32*) and other cytokines such as *TNF*, *CSF*, and *PTGS2* (Figure 3C). IFNγ stimulation alone induced a similar number of cytokines, with reduced intensity and notable exceptions in downregulation of *CCL20*, *CXCL6, IL1A, Il23A, IL33*. IFNλ1 demonstrated the most selective cytokine induction with notable upregulation of *CXCL10, CXCL11, CXCL16*, *CCL22, IL1A, IL1B, IL7*, and *CSF1* and downregulation of *IL23A* (Figure 3C).

Building upon our transcriptomic analysis, we performed comprehensive protein-level validation through multiplex quantification of 40 cytokines in conditioned media from treated IEOs. LMM statistical analysis revealed that the cytokine secretion pattern aligns with our transcriptomic findings. IFNγ and IFNγ + TNF stimulation were potent inducers, significantly regulating 26 and 29 chemokines, respectively, encompassing multiple cytokine families observed in our gene expression analysis. In contrast, IFNλ1 stimulation specifically induced only CXCL10 and CXCL11 secretion, thereby reflecting the more restricted inflammatory signature observed at the transcriptional level (Figure 4A).

**Figure 4:**
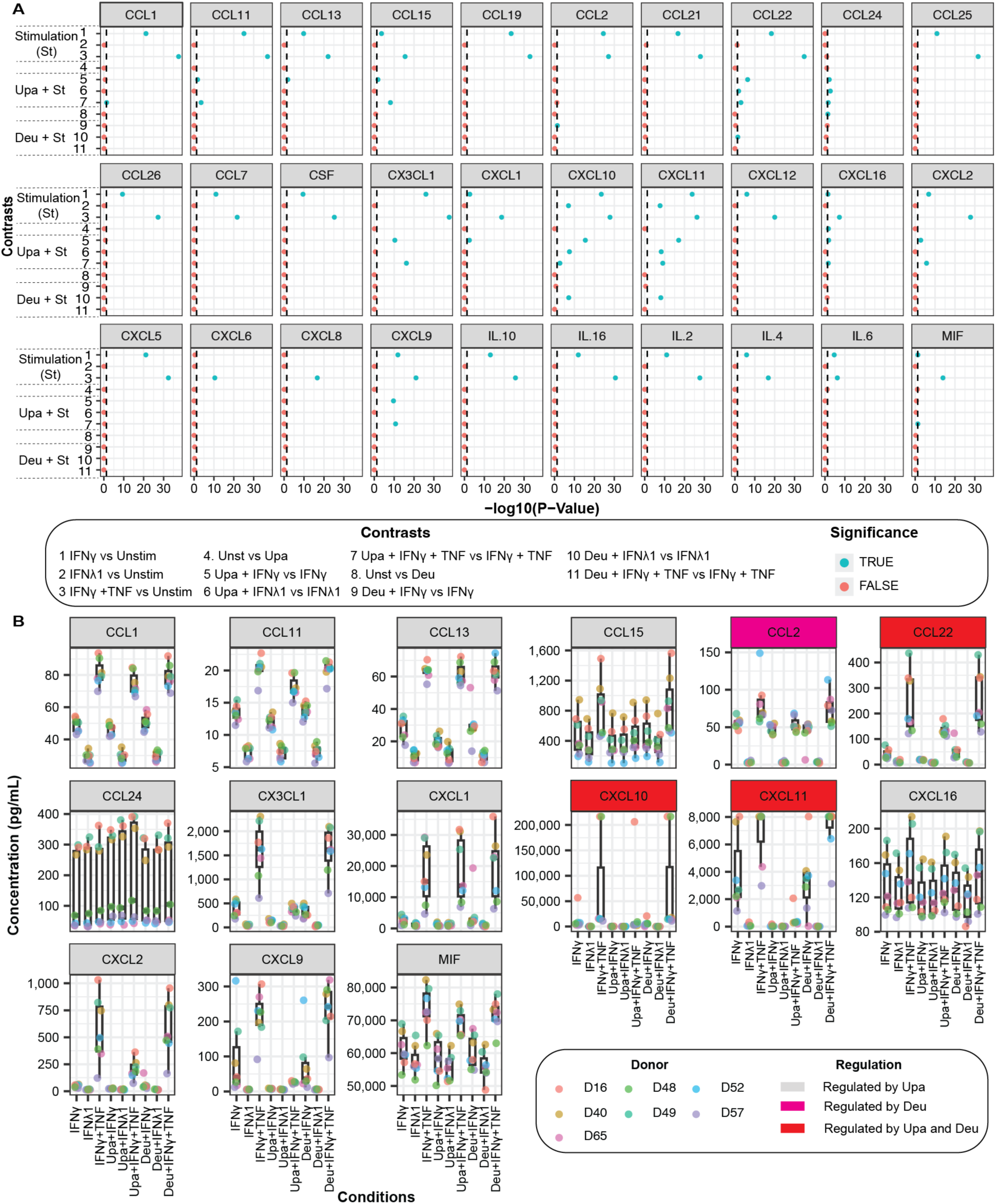
Upadacitinib and deucravacitinib differentially affect chemokine secretion under cytokine-stimulated IEOs. Multiplex detection of chemokines in conditioned medium from IEOs (*n* = 7) in response to stimulation alone or together with inhibitor pretreatment as indicated (1–11). A) Each panel represents a distinct cytokine, with the Y-axis indicating different comparisons and the X-axis representing the negative log10 of adjusted p-values analyzed by linear mixed model (LMM). Statistical significance for the different contrasts is indicated by blue dots above the horizontal dashed line at -log10 (0.05). P-values were adjusted using Benjamini-Hochberg method for multiple comparisons testing. B) Box plots display the concentration (ng/mL) of selected chemokines where at least one drug demonstrated significant regulation. Upa - Upadacitinib, Deu – Deucravacitinib. Raw data are provided in Supplementary File 4.

In IFNs/TNF stimulated IEOs, pretreatment with upadacitinib or deucravacitinib resulted in distinct inhibitory signatures. Upadacitinib demonstrated broad suppression of gene expression across both JAK-STAT signaling and NFκB networks, significantly downregulating transcription factors and downstream effectors in IFNs/TNF stimulated condition. Upadacitinib also significantly inhibited JAK receptor genes and downstream STAT mRNAs (Figure 3A, B). This comprehensive inhibition of JAKs extended to other receptors mRNA expression, with upadacitinib downregulating TNF receptors (*TNFRSF1A, TNFRSF1B*) and multiple interleukin receptors (*ILR1, IL2RG, IL6R, IL6ST, IL7R, IL10RB, IL13RA1, IL15RA, IL17RB, IL18R1, IL18BP, IL22RA1*), while upregulating *IL17RE*, *IL20RA*. Further, upadacitinib pretreatment inhibited expression of cytokine receptors *OSMR, CXCR4, ACKR2* and *ACKR4*, highlighting the integrated nature of these inflammatory networks (Figure 3D). These inhibitory effects further extended to downregulation of CXC, CC, and IL family of cytokines (Figure 3C). Crucially, this transcriptional suppression translated into functional attenuation of secreted inflammatory profiles, with upadacitinib significantly reducing protein levels of 12 out of 26 chemokines that were induced by either IFNγ or IFNγ + TNF stimulation (Figure 4B). This concordance between mRNA and protein-level inhibition underscores the comprehensive immunomodulatory capacity of JAK1-specific inhibition across multiple levels of inflammatory signal transduction. In contrast, deucravacitinib exhibited a more selective and potent inhibition on IFNλ1-induced gene expression, with minimal effects on IFNγ-induced pathways. This TYK2 inhibitor failed to affect IFNγ-induced NFκB signaling components and inhibited *TLR3, MYD88*, and *TNFSF13* (Figure 3A, B). The selective inhibition pattern extended to receptor regulation, with deucravacitinib showing no influence on TNF receptor genes while selectively inhibiting IFNλ1-induced interleukin receptors, *OSMR*, and *ACKR4* and chemokines (Figure 3D). Also, consistent with transcriptomic findings, protein-level analysis revealed that deucravacitinib shows no major impact on secreted chemokines (Figure 4B). These findings highlight the pathway specific role of TYK2 inhibitor in type III interferon signaling.

### Upadacitinib and deucravacitinib attenuates innate immune responsive genes

While previous studies have documented the influence of major histocompatibility complex (MHC-I and II) and antiviral responses in inflammation (33, 78, 79) and how JAK inhibitors affect its expression in IECs (32), the comparative effects of JAK1-specific upadacitinib or TYK2- specific deucravacitinib on MHC regulation or innate antiviral responses remains unexplored. Our transcriptome analysis revealed stimulus-dependent regulatory patterns of MHC-related genes (Figure 3E). MHC-I genes showed consistent upregulation across IFNs/TNF stimulated conditions (except for *HLA-G*). In contrast, MHC-II genes such as *HLA-DMA, HLA-DOB, HLA-DPA1, HLA-DQB1, HLA-DRA, HLA-DRB1, HLA-DRB5* were only regulated between IFNγ and IFNγ + TNF-stimulated conditions. Pretreatment with upadacitinib downregulated expression of IFNγ, IFNλ1, and IFNγ + TNF-induced MHC-I and MHC-II genes. In contrast, deucravacitinib only had a significant effect on IFNλ1-induced MHC-I, while no effect on any IFNs/TNF stimulated MHC-II expression (Figure 3E).

Extending our analysis to pattern recognition receptor expression and antiviral response genes, we characterized the transcriptional regulation of genes encoding viral sensors, signal transducers, and effector molecules in IEOs stimulated with IFNγ, IFNλ1 and IFNγ + TNF (Figure 3F). IFNs/TNF stimulation induced a strong activation of ISGs, with key activation of interferon-induced protein with tetratricopeptide repeats family genes (*IFIT1, IFIT2, IFIT3*), interferon-induced transmembrane proteins (*IFITM1, IFITM2, IFITM3*), and myxovirus resistance proteins (*MX1, MX2*). Critical sensors of viral nucleic acids (*CGAS, STING1, IFIH1/RIG1*, *DHX58, IFI16, ZBP1*) and antiviral effector molecules (*TBK1, IRF3, ISG15, ISG20, OASL, OAS1, OAS2, OAS3*) were similarly upregulated. The APOBEC3 family of cytidine deaminases were also significantly induced, complemented with strong expression of innate immune effectors including guanylate-binding proteins (*GBP1, GBP2, GBP3*) and TRIM family members. Pretreatment with upadacitinib effectively downregulated expression of IFNγ, IFNλ1 and IFNγ + TNF-induced ISGs, innate immune effectors and cytosolic DNA sensors (Figure 3F). In contrast, deucravacitinib demonstrated potent and selective inhibition of IFNλ1-induced gene expression and exhibited inhibition of only a subset of genes induced by IFNγ + TNF stimulation.

### Upadacitinib attenuates cell death-related pathways and promotes cell cycle, DNA replication, and developmental signaling programs downregulated in IFNs/TNF-stimulated IEOs

IECs respond to inflammatory cytokines, which not only activate immune responses but also substantially influence metabolic stress (80), epithelial barrier function (40) and various forms of cell death (37, 81), thereby disrupting epithelial homeostasis. Previous studies have reported that chronic inflammation leads to dysregulation of cell cycle regulators, particularly cyclin-dependent kinases, and key developmental signaling such as WNT and Hedgehog (82–85). These alterations impair epithelial renewal and can contribute to increased cancer risk in IBD patients (86, 87). Given this context, it is critical to understand how inflammatory cytokines modulate the balance between epithelial cell death and stemness and how selective inhibition of JAK1 or TYK2 may restore these networks to promote mucosal healing in epithelial cells. To elucidate this, we examined the transcriptional regulation of cell death, cell cycle, DNA replication, and developmental signaling in IEOs under IFNs/TNF stimulation and assessed the differential impact of JAK1 and TYK2-specific inhibition on these interconnected processes. Over-representation enrichment analysis revealed that IFNs/TNF-stimulated conditions profoundly induced cell death pathways while suppressing proliferative and developmental pathways (Supplementary Figure 5, 6). Transcriptional analysis of cell death pathways demonstrated stimulus-specific induction patterns. IFNγ and IFNγ + TNF stimulation strongly upregulated expression of extrinsic and intrinsic apoptotic factors (*NOD2, BAX1, BAK1, BBC3, CFLAR, CARD9, CARD14, TRADD, FAS, TNFSF10, XAF1*) culminating in activation of executioner caspases (*CASP3, CASP7*, and *CASP10*) (Figure 5A). In contrast, IFNλ1 stimulation elicited minimal apoptotic gene induction limited to *BAK1, BBC3, CASP7*, *CASP8, CASP10, CFLAR, FAS, NOD2, XAF1, TNFSF12* (Figure 5A). Furthermore, all IFNs/TNF stimulation strongly upregulated pyroptosis-associated genes (*CASP1, GSDMD*) (Figure 5C), necroptosis genes (*RIP, CYLD, MLKL*), PANoptosis genes (*ZBP1, CASP8*) (Figure 5D), NADPH oxidase components (*NOX1, DUOX2, DUOXA2*) and nitric oxide synthase (*NOS2*) (Figure 5E). Notably, IFNγ and IFNγ + TNF stimulation exhibited stimulus-specific induction of antioxidant defense gene (*SOD2, GPX2*) and downregulated multiple cytoprotective factors (*FOS, GLRX2, GPX4, HMOX1, NQO1, UCP2*) (Figure 5E).

**Figure 5:**
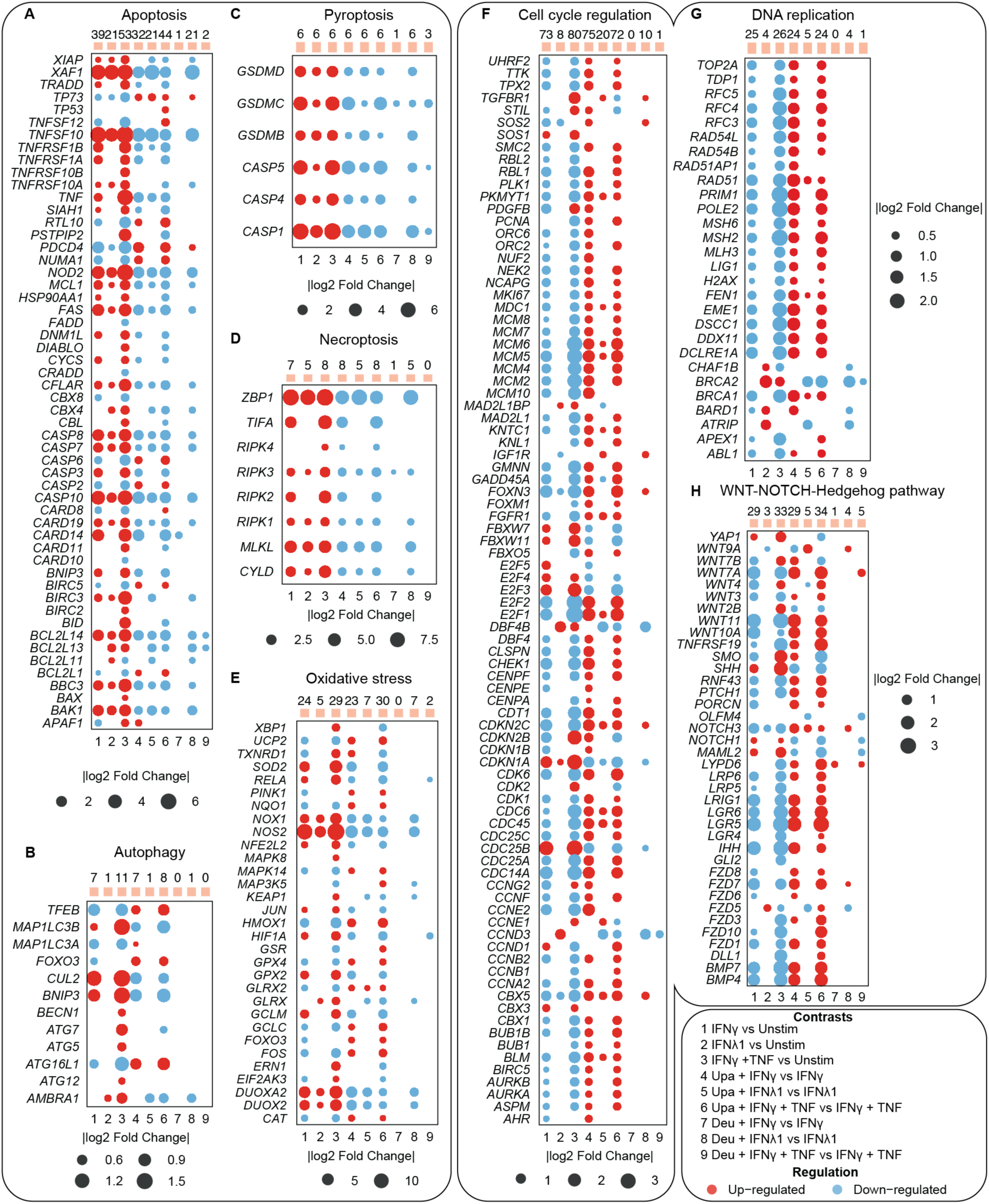
Upadacitinib and deucravacitinib distinctly regulate cell death and cell-cycle gene expression cytokine-stimulated IEOs. Bubble plots showing log2 fold change of selected genes in A) Apoptosis, B) Autophagy C) Pyroptosis, D) Necroptosis, E) oxidative stress, F) Cell cycle regulation, G) DNA replication, and H) WNT/NOTCH/Hedgehog signaling in IEOs (*n* = 7) stimulated with IFNγ, IFNλ1, and IFNγ + TNF (1–3) alone or together with pretreatment of upadacitinib (Upa) (4–6) or deucravacitinib (Deu) (7–9). Red and blue circles indicate up- and down-regulation, respectively, with circle size proportional to fold change. Only genes with p-value p < 0.05 are presented.

The convergence of multiple cell death pathways (Figure 5A-E) under IFNγ + TNF stimulation suggests a mechanistic framework for understanding extensive epithelial damage in severe inflammatory condition. Pretreatment of upadacitinib comprehensively downregulated genes associated with different cell death pathways across IFNs/TNF stimulation conditions, while deucravacitinib exhibited selective modulating patterns predominantly affecting IFNλ1-induced gene expression (Figure 5A-E) compared to IFNγ/TNF-induced cell death programs, with distinct dependencies on JAK versus TYK2-mediatied signaling (Figure 5A-E).

Our transcriptional analysis on cell cycle related pathways revealed that IFNγ and IFNγ + TNF conditions substantially downregulated cyclin-dependent kinases (*CDK1, CDK4, CDK6*), their regulatory cyclins (*CCNA2, CCNB1, CCNB2*), cell cycle phosphatases (*CDC25A, CDC25C*), mitotic checkpoint components (*BUB1, BUB1B, MAD2L1*) and the chromosome segregation machinery (*CENPA, CENPE, CENPF*) (Figure 5F). This cell cycle arrest signature encompassed transcriptional regulators with more pronounced downregulation of E2F factors (*E2F1, E2F2*), and *FOXM1*, a master regulator of G2/M progression. IFNγ + TNF exhibited selective upregulation of cell cyclin mediators as well (*CDK2, CDK7, CDK9, E2F3, E2F4 CDKN22B, CDC25B*) (Figure 5F). This synchronized regulation extended to DNA replication, where IFNγ and IFNγ + TNF-stimulated conditions downregulated origin recognition complex components (*ORC2, ORC6*), DNA licensing factors (*CDC6, CDT1*), the MCM helicase complex (*MCM2-MCM7*), core replication factors (*POLE2, PRIM1, RFC3-RFC5, MKI67, PCNA*) and processing enzymes (*FEN1, LIG1*) (Figure 5G). Similarly, DNA damage response pathways exhibited coordinated downregulation of DNA repair elements (*RAD51, RAD51AP1, RAD54AB, RAD54L, BRCA1, MSH2, MSH6*) (Figure 5G).

The integrated nature of these cellular programs was further investigated by looking into developmental signaling networks which are critical for intestinal epithelial homeostasis and stem cell maintenance (Figure 5H). IFNγ and IFNγ + TNF- stimulated conditions showed substantial downregulation of canonical WNT receptors and ligands (*LGR4/5/6, LRP5/6, WNT3, WNT7A*), and Frizzled receptor expression (*FZD1, FZD3, FZD5, FZD7, FZD8, FZD10*), *NOTCH* signaling pathway component (*NOTCH3*), Hedgehog pathway elements (*IHH, GLI2, PTCH1*) (Figure 5H). In contrast, IFNγ + TNF selectively upregulated specific WNT receptors (*WNT2B, WNT4, WNT7B*), NOTCH elements (*NOTCH1, MAML2*) and hedgehog regulators (*SHH, SMO*). Notably, intestinal stem cell markers such as *LGR5* and *TNFRSF19* showed pronounced downregulation following both IFNγ and IFNγ + TNF stimulation indicating reduced regenerative capacity under inflammatory conditions (Figure 5H). In general, IFNγ and IFNγ + TNF stimulations were significantly enriched for downregulation of cell cycle related process networks compared to IFNλ1 stimulation (Supplementary Figure 5, 6).

Pretreatment with upadacitinib normalized these interconnected transcriptional networks. For example, upadacitinib significantly reversed IFNγ and IFNγ + TNF-mediated suppression of cell cycle regulators (*CDK1, CDK6, CCNA2, CCNB1, CCNB2, CDC25A, CDC25C, BUB1, BUB1B, MAD2L1, CENPA, CENPE, CENPF, E2F1, E2F2, FOXM1)*, while downregulating IFNs/TNF induced cell cycle modulators (*CDK2, CDK7, CDKN22B, CDC25B*) (Figure 5F). This coordinated upregulation extended to DNA replication (*ORC2, CDC6, CDT1, MCM2-MCM7, POLE2, PRIM1, RFC3-RFC5, MKI67, PCNA, FEN1, LIG1*) and repair elements (*RAD51, RAD51AP1, RAD54AB, RAD54L, BRACA1, MSH2, MSH6*) maintaining genomic integrity networks critical for epithelial renewal (Figure 5G). Also, upadacitinib pretreatment normalized developmental signaling pathways by upregulating canonical WNT receptors and ligands (*LGR4/5/6, LRP5/6, WNT3, WNT7A*), Frizzled receptors (*FZD1, FZD3, FZD5, FZD7, FZD8, FZD10*), NOTCH components (*NOTCH3*), and Hedgehog elements (*IHH, GLI2, PTCH1*) reduced by IFNs/TNF, while downregulating IFNs/TNF-induced factors (*WNT2B, WNT 7B, NOTCH1, MAML2, SHH, SMO*) (Figure 5H). The restoration of intestinal stem cell markers (*LGR5, TNFRSF19*) further demonstrated the potential of upadacitinib pretreatment to restore mucosal healing capacity under inflammatory conditions. In contrast, deucravacitinib exhibited no effect on IFNγ and IFNγ + TNF-induced regulation of gene expression related to proliferation and developmental signaling (Figure 5F-H). These findings show that upadacitinib may directly promote mucosal healing beyond conventional anti-inflammatory effects.

### JAK inhibitor treatment is associated with elevated colonic epithelial proliferation *in vivo*

Because Ki67 is expressed throughout all active phases of the cell cycle (G1, S, G2, M) but absent in quiescent cells (G0) (88), it serves as a reliable marker to investigate the regulation of cell cycle and DNA replication programs identified in our transcriptomic analysis. We therefore assessed epithelial proliferation by Ki67 immunohistochemistry in colonic biopsies from healthy controls (HC, *n* = 10) and two comparable groups of UC patients mainly on no medication (*n* = 14) or treated with JAK inhibitors (filgotinib or tofacitinib) (Table 2). The percentage of Ki67+ epithelial cells was significantly elevated in JAK inhibitor-treated patients compared to both HC and UC controls (Figure 6, Supplementary File 3; unpaired Wilcoxon rank-sum tests with Benjamini-Hochberg correction, all FDR ≤ 0.001), and this effect was observed regardless of histological inflammation status. As one of the JAK inhibitor-treated patients also received 5-ASA, we included one 5-ASA-treated patient in the comparing UC group (both marked with black circle in Figure 6). The epithelial Ki67 proliferation index of these two patients did not stand out from the rest and excluding them from the analyses did not alter the result. These *in vivo* findings in patients on filgotinib or tofacitinib support the transcriptional restoration of cell cycle and DNA replication programs observed under upadacitinib in our IEO model.

**Figure 6:**
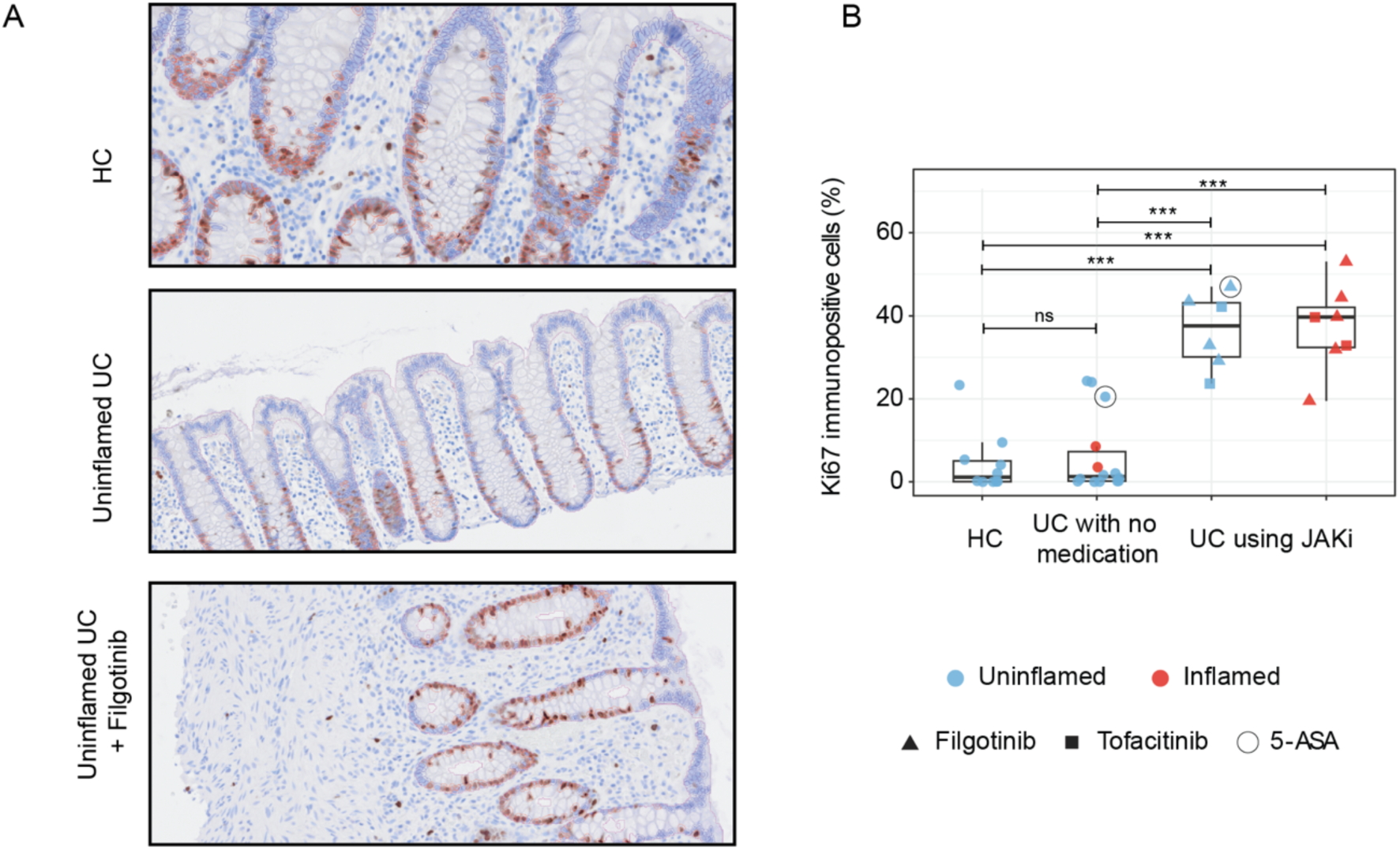
Ki67 is significantly increased in patients taking JAK inhibitors. Sections of colonic mucosa of healthy control (HC), UC patients mainly on no medication and patients under JAK inhibitor were stained for Ki67 by immunohistochemistry and analyzed in QuPath **A)** Representative images of segmented epithelium with Ki67+ cells (red circles) are shown from HC, UC patients mainly on no medication and weakly inflamed patient under filgotinib. B) Box plots show the percentage of Ki67+ cells per patients, across three groups: HC (*n* = 10), UC patients with no medication (uninf-UC, *n* = 14), and UC patients using JAK inhibitors (uninflamed, *n* = 6; inflamed, *n* =7). Differences between groups were assessed using unpaired Wilcoxon rank-sum tests (Mann-Whitney U) with Benjamini-Hochberg correction across the specified comparisons. *FDR < 0.05, **FDR < 0.01, ***FDR < 0.001; ns = not significant.

## Discussion

A central finding of our study is that JAK/TYK2 inhibitors exert a strong impact on IFNs/TNF-induced genes associated with JAK-STAT and NFκB signaling cascades, modulation of innate antiviral defense, attenuation of cell death pathways and finally regulation of cell cycle, DNA replication, and developmental signaling processes which are involved in induction of mucosal healing (9, 12). IEOs are increasingly used as a tool in *ex vivo* drug screening and therapeutic testing, for evaluating drug efficacy, and toxicity in patient-specific manner (89). However, previous studies of JAK inhibitors effects on IEOs have largely focused on cytokine-induced barrier dysfunction, cell death or cytokine release (32, 34, 37, 38, 40, 90–93), and have not systematically examined how these drugs influence epithelial response at the transcriptional level in a cytokine specific context. Our study addresses this by evaluating the receptor-specific transcriptional response to JAK/TYK2 inhibitors at a defined time point (8 hours post cytokine stimulation following 16 hours of drug pretreatment). For instance, under IFNγ/TNF stimulation, upadacitinib excerpted broader transcriptional regulation of inflammatory pathways compared to deucravacitinib. IFNλ1-induced gene expression was strongly modulated by both inhibitors, while TNF-driven transcriptional changes were minimally affected consistent with TNF signaling through a JAK-independent mechanism. To our knowledge, this is the first study that provides insights into the multifaceted roles of JAK inhibition in modulating intestinal epithelial responses to interferons and TNF, extending beyond their established role in immune cell modulation in IBD.

Inhibitors with different JAK selectivity are expected to exert distinct cellular effects depending on the receptor-ligand context (19, 30), providing a mechanistic basis for the different transcriptional responses observed. Our comparative analysis revealed clear differences with JAK inhibitors (upadacitinib > brepocitinib > tofacitinib > filgotinib) significantly inhibited interferon-induced STAT1 and STAT3 phosphorylation, while deucravacitinib inhibited TYK2 phosphorylation in a dose dependent manner. Few studies have investigated the effects of JAK inhibitors on cytokine-induced phosphorylation of STAT3 and TYK2 in colonic epithelial organoids, but inhibition of STAT1 phosphorylation has been reported (32, 38, 90).

These distinct phosphorylation patterns provide a mechanistic basis for the divergent downstream effects. The intestinal epithelium maintains a sophisticated antiviral defense system that relies heavily on interferon signaling (23, 79) and previous findings highlighted the essential role of interferons, particularly IFNλ, in constraining multiple intestinal viral infections (94, 95). Constant et al. (24), reported that type III IFNs were least cytotoxic compared to type II, suggesting a protective role in maintaining epithelial integrity during antiviral response. Our observed differential effects of upadacitinib and deucravacitinib on antiviral responses carry significant clinical implications. Both inhibitors substantially suppressed antiviral responses in IFNs/TNF-stimulated conditions, while only deucravacitinib specifically suppressed IFNλ1-induced responses. These findings align with a previous *in vitro* study on Caco-2 cell line, which reported that both baricitinib (JAK1/2) and deucravacitinib (TYK2) inhibited IFNα and IFNλ1-induced antiviral responses (95). However, the study found that compared to baricitinib, deucravacitinib (at 0.25 μM) was less effective in inhibiting IFNλ1-induced antiviral responses than IFNα-induced expression. The authors conferred that at clinically relevant concentration, a TYK2 inhibitor might well preserve type III (IFNλ) induced antiviral responses compared to broader acting JAK1/2 inhibitor in IECs (95) and might potentially translate to reduced infection risk in the clinical practice. Also, the broad inhibitory effect of upadacitinib on antiviral genes aligns with clinical observation of increased infection risk (96).

Cell death is a critical determinant of epithelial turnover and mucosal inflammation in IBD. Our transcriptome analysis revealed that IFNγ, IFNλ1, IFNγ + TNF stimulation of IEOs recapitulated cell death signatures observed in UC patients (37, 97–100). Our data indicate a more prominent role of necroptotic, and pyroptotic cell death mechanisms compared to apoptosis at the concentration used in our study (IFNγ at 10 ng/mL + TNF at 50 ng/mL), whereas higher doses of IFNγ + TNF (50 ng/mL each) have been reported to favor mitochondrial apoptosis alongside contribution from necroptotic or pyroptosis (37). Consistent with the inhibitor-specific patterns observed for antiviral responses, upadacitinib broadly attenuated IFNs/TNF-induced cell death programs. Deucravacitinib on the other hand displayed selective modulation, predominantly affecting IFNλ1-induced cell death with minimal effects on IFNγ + TNF-induced responses. Consistent with our findings, the capacity of JAK inhibitors to mitigate cytokine-induced cell death in IECs has been previously documented in UC and CD colonoids (97) and in CD enteroids (36). Mechanistically, upadacitinib-mediated suppression of IFNs/TNF-induced NOS2 expression aligns with the proposed regulation of PANoptosis via the JAK-STAT-IRF1-iNOS-NO pathway (101). These findings extend previous observation on JAK inhibitors effects on IFNγ (50 ng/mL) induced iNOS expression (90), highlight the molecular mechanism by which the JAK inhibitors may attenuate inflammatory epithelial damage.

Beyond suppressing cell death, JAK inhibition also influenced epithelial proliferation. Cytokines exert opposing effects on intestinal epithelial proliferation (102). IL22 has shown to drive epithelial proliferation via STAT3 activation (103), while IFNγ, essentially in combination with TNF, inhibits stem cell proliferation and increases apoptosis (104). Consistent with this, our transcriptional analysis also revealed that IFNγ and IFNγ + TNF conditions substantially downregulated genes involved in cell cycle/DNA replication related pathways that upadacitinib reversed this suppression. Clinically, the OCTAVE trial of tofacitinib (25) and the SELECTION trial of filgotinib (27) have established endoscopic and mucosal healing as primary endpoints, but the cellular mechanism underlying this healing has not been directly addressed, and to our knowledge no study has directly demonstrated whether JAK inhibition drives epithelial proliferation in patients. Despite the small sample size, our Ki67 data shows that UC patients on JAK inhibitors (*n* = 11, 13 biopsies) have enhanced epithelial proliferation *in vivo* and this could mechanistically be due to the reversal of IFNγ-mediated proliferative arrest. Notably, elevated Ki67 was observed in JAK inhibitors treated UC patients regardless of mucosal inflammation status, suggesting a direct effect of JAK inhibition on epithelial proliferation. Whether sustained epithelial proliferation under long-term JAK inhibition carries oncogenic implications warrants further investigation. According to the most recent ECCO guidelines, there is currently no evidence of increased overall cancer risk in patients with IBD treated with JAK inhibitors, although the available evidence is derived from randomized controlled trials and long-term data in IBD remain limited (105). Whether the proliferative response we observe represents physiological restoration of homeostatic regeneration or pharmacologically driven hyperproliferation cannot be determined from Ki67 alone and will require follow-up studies including assessment of crypt architecture, stem cell markers (e.g., LGR5), and surveillance for dysplastic features in patients on long-term therapy.

A key strength of this study is the comparative analysis of multiple JAK inhibitors in colonoids derived from UC patients, providing unique insights into the differential effects on epithelial responses relevant for antiviral, cell death and mucosal healing in the intestinal epithelium. A limitation of our study is that we use a subset of cytokines to model the complex inflammatory milieu, and a relatively short stimulation period which may not fully capture the chronic effects of inflammatory cytokines in intestinal epithelial cells. Although we did not perform time-course transcriptome experiments, our approach allowed us to capture early transcriptional signatures that reflect the immediate impact of JAK inhibition on cytokine signaling. Further, *in vivo* validation of other genes promoting mucosal healing using relevant patient material is needed to strengthen the translational relevance of our findings.

## Conclusions

In summary, our comprehensive analysis of JAK/TYK2 inhibitors effect on patient-derived IEOs challenges the current paradigm of JAK inhibitors as solely immune-modulatory agents and reveals their capacity to directly promote genes involved in epithelial repair and restoration. By elucidating how different JAK inhibitors differentially regulate epithelial responses to inflammatory stimuli, our study provides insights for optimizing therapeutic efficacy while minimizing adverse effects, ultimately advancing toward personalized JAK inhibitor therapy that addresses both immune dysfunction and epithelial restoration in UC patients.

## List of abbreviations

BEST4: Bestrophin 4
CCL: Chemokine (C-C motif) ligand
CXCL: Chemokine (C-X-C motif) ligand
DEG: Differentially expressed gene
DUOX2: Dual oxidase 2
DMSO: Dimethyl sulfoxide
EGFR: Epidermal growth factor receptor
GAPDH: Glyceraldehyde 3-phosphate dehydrogenase
GAS: Gamma-activated sequence
HC: Healthy control
IBD: Inflammatory bowel disease
IEC: Intestinal epithelial cell
IEO: Intestinal epithelial organoid
IFN: Interferon
IL: Interleukin
iNOS: inducible nitric oxide synthase
IRF: Interferon regulatory factor
ISG: IFN-stimulated genes
ISREs: Interferon-stimulated response elements
JAK: Janus kinase
LGR: Leucine-rich G repeat-containing protein-coupled receptor
LMM: Linear mixed model
log2FC: Log2 fold change
MAPK: Mitogen-activated protein kinase
MHC: Major histocompatibility complex
MLCK: Myosin light chain kinase
mRNA: Messenger ribonucleic acid
NFκB: Nuclear factor kappa-light-chain-enhancer of activated B ells
OTOP2: Otopetrin 2
PI3K: Phosphoinositide 3-kinase
PROGENy: Pathway RespOnsive GENes
RIPK: Receptor-interacting protein kinases
S1P: Sphingosise-1-phosphate
Seq: Sequencing
STAT: Signal transducer and activator of transcription
TNF: Tumor necrosis factor
TYK2: Tyrosine kinase 2
UC: Ulcerative colitis
WB: Western blot
WNT: Wingless-related integration site

## Declarations

### Ethics approval and consent to participate

Use of patient-derived IEOs and *in vivo* samples from Trondheim was approved by the Central Norway Regional Committee for Medical and Health Research Ethics, reference numbers 22687 and 26789. The *in vivo* samples from Oslo were obtained from the OUS-IBD biobank (reference number 64905) and approved for use in the current study reference number 860752. All patients gave written informed consent. All experimental methods were performed adhering to the principles of the Declaration of Helsinki.

### Consent for publication

Not applicable.

### Availability of data and materials

The RNA sequencing datasets generated and analyzed during the current study are available in the Array express and are accessible through accession numbers E-MTAB-16024. All supplementary files (1–6), including associated metadata, used for figure generation, are under supplementary material. Any additional information required to reanalyze the data reported in this paper is available from the lead contact upon request.

### Authors contributions

AS contributed to this work as a first authors. AS, MDH, IB, and TB conceptualized the study. AS, GAW, MDH, SS, JPM, RC, MGM, LG, IB and TB conducted methodology and the validation, while GAEW and AS performed the software programming and formal analysis. Investigation was performed by AS, GAW, SS, JPM, RC, MGM, and LG. Resources were provided by MDH, MO, MLH, AEØ, IB, and TB. Data curation was managed by AS, IB, and TB. The original draft of the manuscript was written by AS, IB, and TB. All authors reviewed and edited the manuscript and approved the final version. AS created the Visualization. MDH, IB and TB supervised the project. Funding acquisition and project administration were undertaken by AEØ, IB, and TB.

## Acknowledgements

This work was performed in collaboration with the Gastrointestinal Endoscopy Unit at the Department of Gastroenterology and Hepatology, St. Olav’s University Hospital, and we thank Kari Lillebråten and Hanne Fossen for their invaluable assistance related to biobanks and IBD patients that were essential for this research. We thank Henrik P. Sahlin Pettersen at the Department of Pathology, St. Olav’s University Hospital, for the epithelial segmentation of immunohistochemistry-stained sections. We also thank Linn Karina M. Selvik at the Department of Clinical and Molecular Medicine (IKOM) for her technical assistance. RNA library preparation, sequencing, parts of the bioinformatics analysis, and cell deconvolution were performed in close collaboration with Vidar Beisvåg, Arnar Flatberg and Tone Christensen at the Genomics Core Facility (GCF), Norwegian University of Science and Technology (NTNU) for immunostaining and scanning of sections. Both GCF and CMIC are funded by the Faculty of Medicine and Health Sciences at NTNU, and the Central Norway Regional Health Authority.

## Declaration of generative AI and AI-assisted technologies

During the preparation of this work, the author(s) used M365 Copilot to edit the text since English is not the authors native language and Claude.ai (Anthropic, San Francisco, CA, USA) was used to aid with writing R syntax. After using this tool or service, the authors reviewed and edited the content as needed and take full responsibility for the content of the publication.

## Corresponding author

Requests for further information and resources should be directed to and will be fulfilled by the lead contact, Torunn Bruland.

## Supplementary figures and legends

**Supplementary Figure 1:**
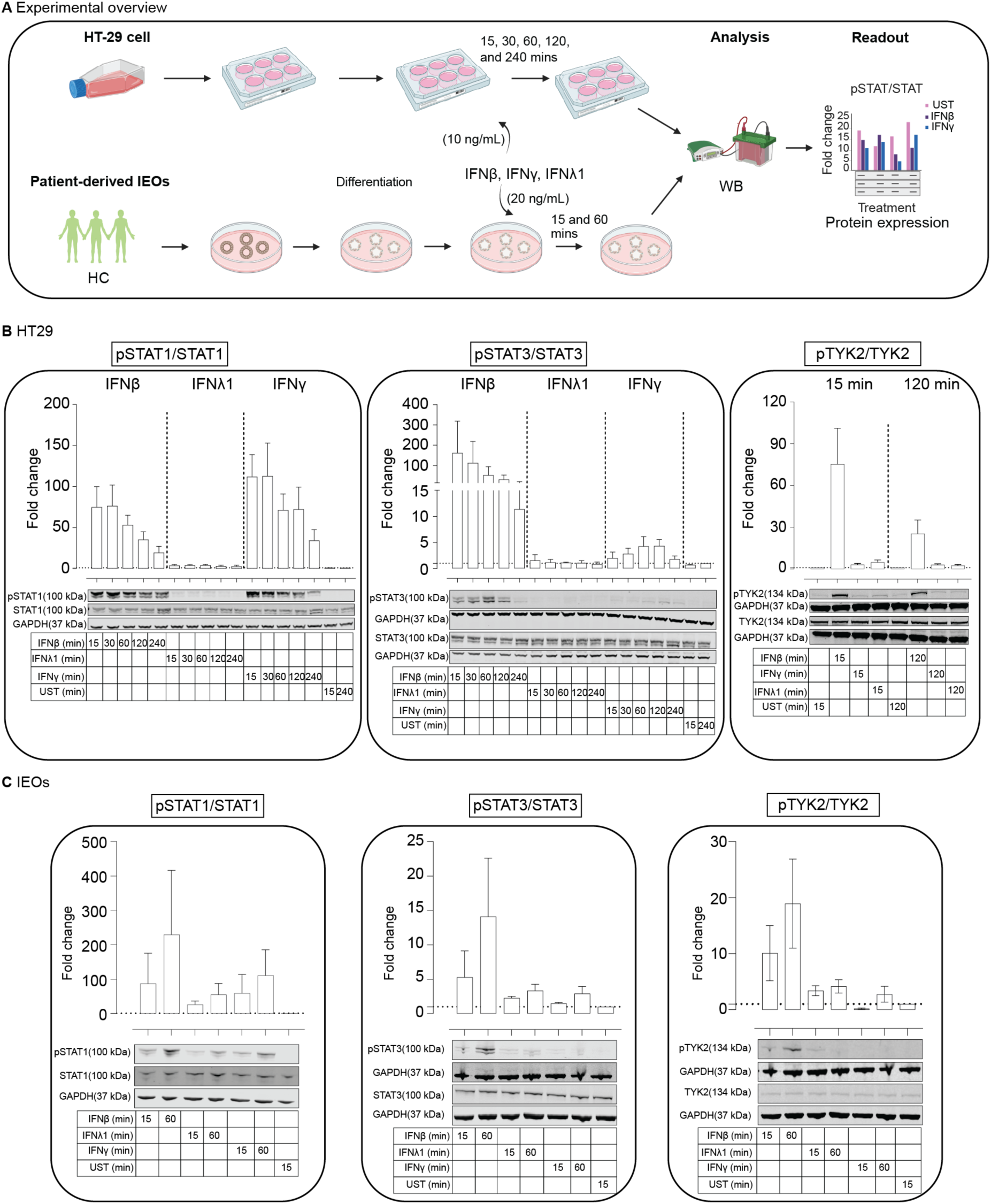
Time series of IFNβ, IFNγ, and IFNλ1-induced activation of pSTAT1, pSTAT3, and pTYK2: A) Schematic representation of experimental overview used in HT-29 cells and patient-derived IEOs. Quantification of phosphorylated STAT1, STAT3, and TYK2 represented as pSTAT1/STAT1, pSTAT3/STAT3, and pTYK2/TYK2 in B) HT29 cells and **C)** IEOs. Fold change is generated by normalizing to unstimulated control (DMSO) and further to GAPDH. Immunoblot for one representative replicate is shown under the quantification, and other replicates are in the Supplementary Figure 10, 11.

**Supplementary Figure 2:**
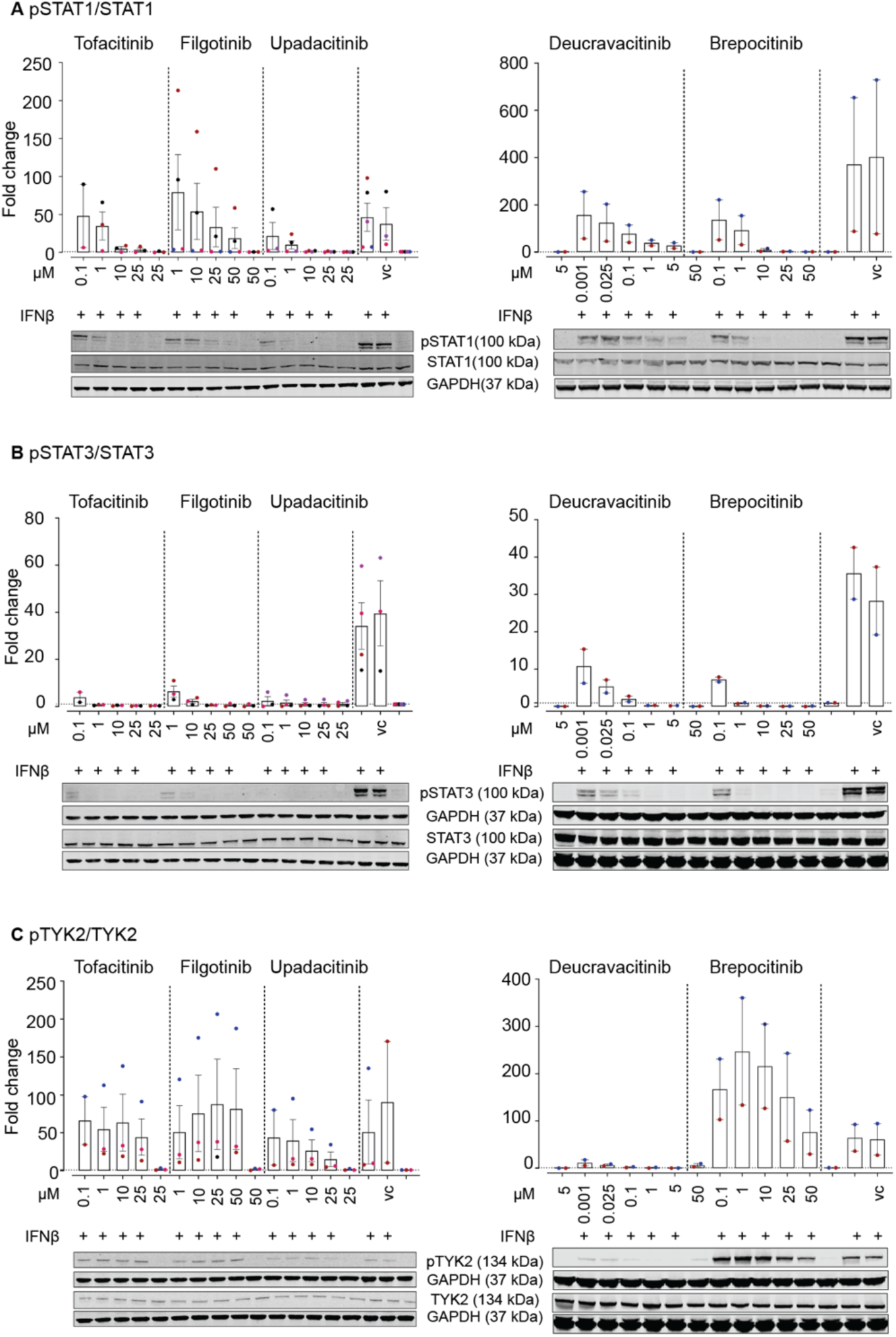
Effects of five JAK-TYK2 inhibitors on IFNβ-induced pSTAT1, pSTAT3, and pTYK2 levels assessed by immunoblotting. (A-C) The bar graphs represent immunoblotting quantification of A) pSTAT1, B) pSTAT3, and C) pTYK2 in IEOs (*n* = 2-5) in response to either DMSO (VC: vehicle control) or tofacitinib (0.1-25 µM), upadacitinib (0.1-25 µM) and filgotinib (1-50 µM), and deucravacitinib (0.001-5 µM), and brepocitinib (0.1-50 µM) for one hour pretreatment with or without IFNβ (20 ng/mL) stimulation for one hour. Fold change of pSTAT1/3 and pTYK2 is calculated by normalization to unstimulated control (negative control) and then to GAPDH. Immunoblot for one representative replicate is shown (other replicates in Supplementary Figure 12, 15).

**Supplementary Figure 3:**
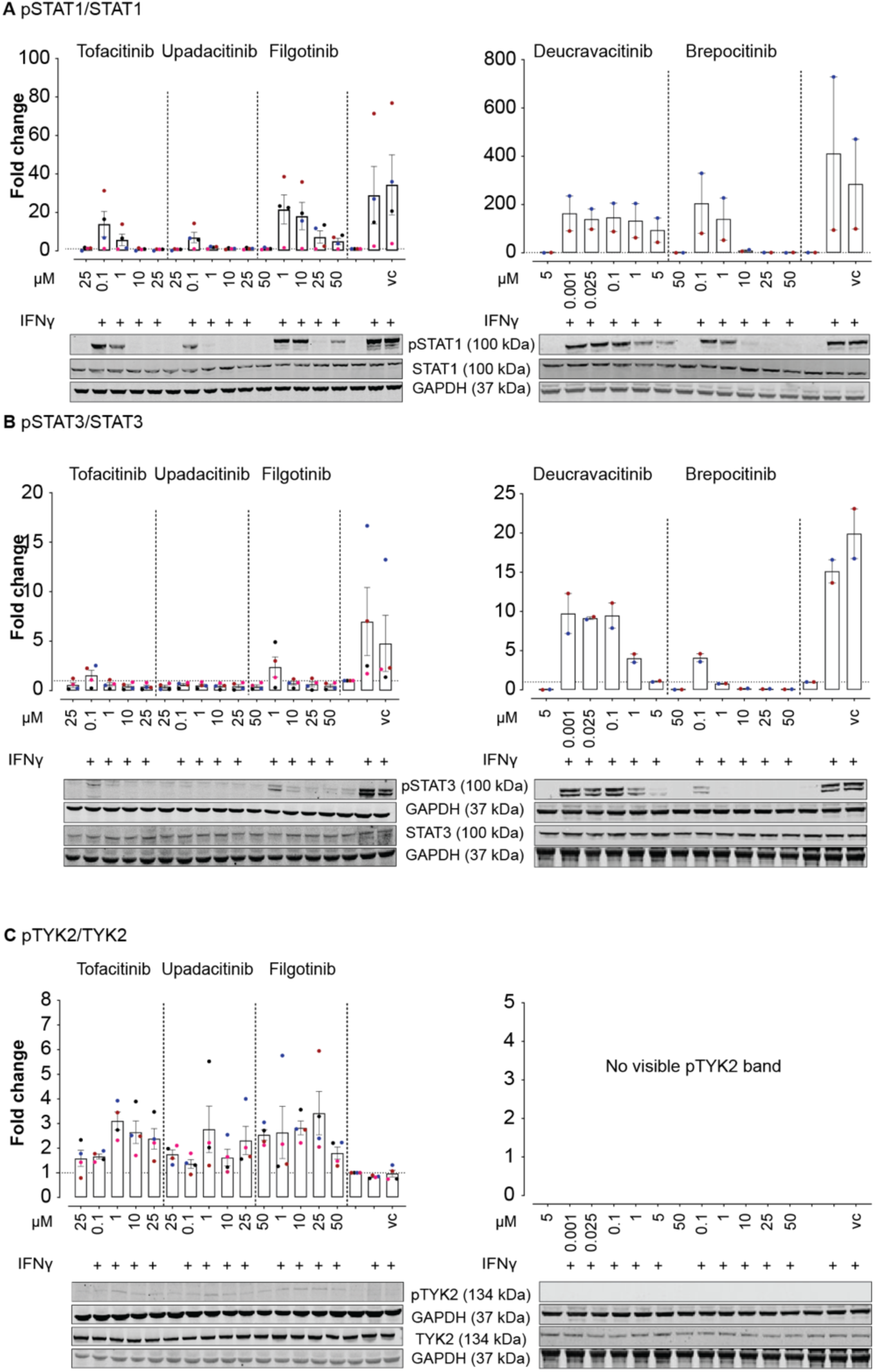
Effects of five JAK-TYK2 inhibitors on IFNγ-induced pSTAT1, pSTAT3, and pTYK2 levels assessed by immunoblotting. (A-C left and middle) The bar graphs represent immunoblotting quantification of A) pSTAT1, B) pSTAT3, and C) pTYK2 in IEOs (*n* = 2-5) in response to either DMSO (VC: vehicle control) or tofacitinib (0.1-25 µM), upadacitinib (0.1-25 µM) and filgotinib (1-50 µM), and deucravacitinib (0.001-5 µM), and brepocitinib (0.1-50 µM) for one hour pretreatment with or without IFNγ (20 ng/mL) stimulation for one hour. Fold change of pSTAT1/3 and pTYK2 is calculated by normalization to unstimulated control (negative control) and then to GAPDH. Immunoblot for one representative replicate is shown (other replicates in Supplementary Figure 13, 15).

**Supplementary Figure 4:**
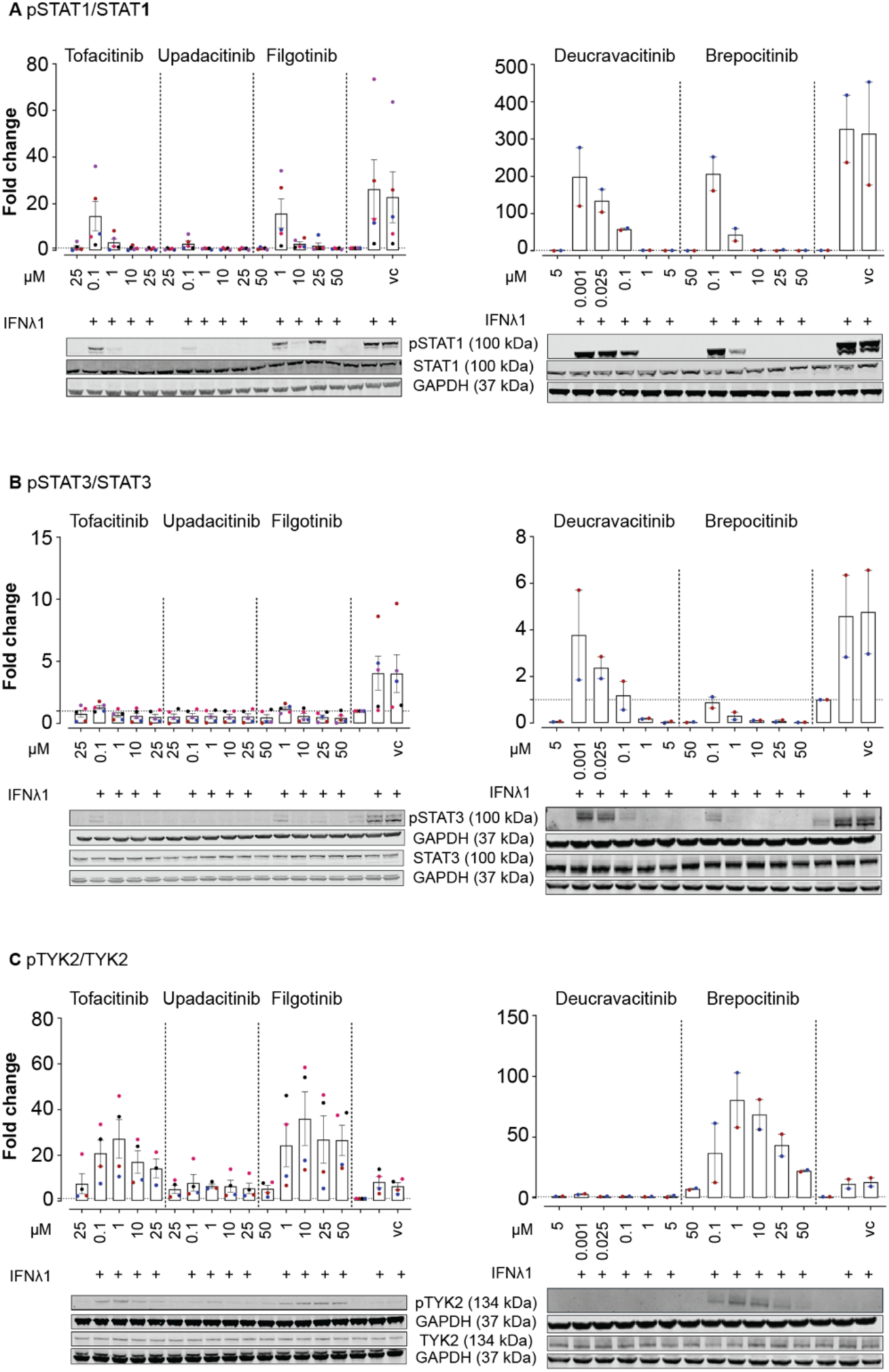
Effects of five JAK-TYK2 inhibitors on IFNλ1-induced pSTAT1, pSTAT3, and pTYK2 levels assessed by immunoblotting. (A-C left and middle) The bar graphs represent immunoblotting quantification of A) pSTAT1, B) pSTAT3, and C) pTYK2 in IEOs (*n* = 2-5) in response to either DMSO (VC: vehicle control) or tofacitinib (0.1-25 µM), upadacitinib (0.1-25 µM) and filgotinib (1-50 µM), and deucravacitinib (0.001-5 µM), and brepocitinib (0.1-50 µM) for one hour pretreatment with or without IFNλ1 (20 ng/mL) stimulation for one hour. Fold change of pSTAT1/3 and pTYK2 is calculated by normalization to unstimulated control (negative control) and then to GAPDH. Immunoblot for one representative replicate is shown (other replicates in Supplementary Figure 14, 15).

**Supplementary Figure 5:**
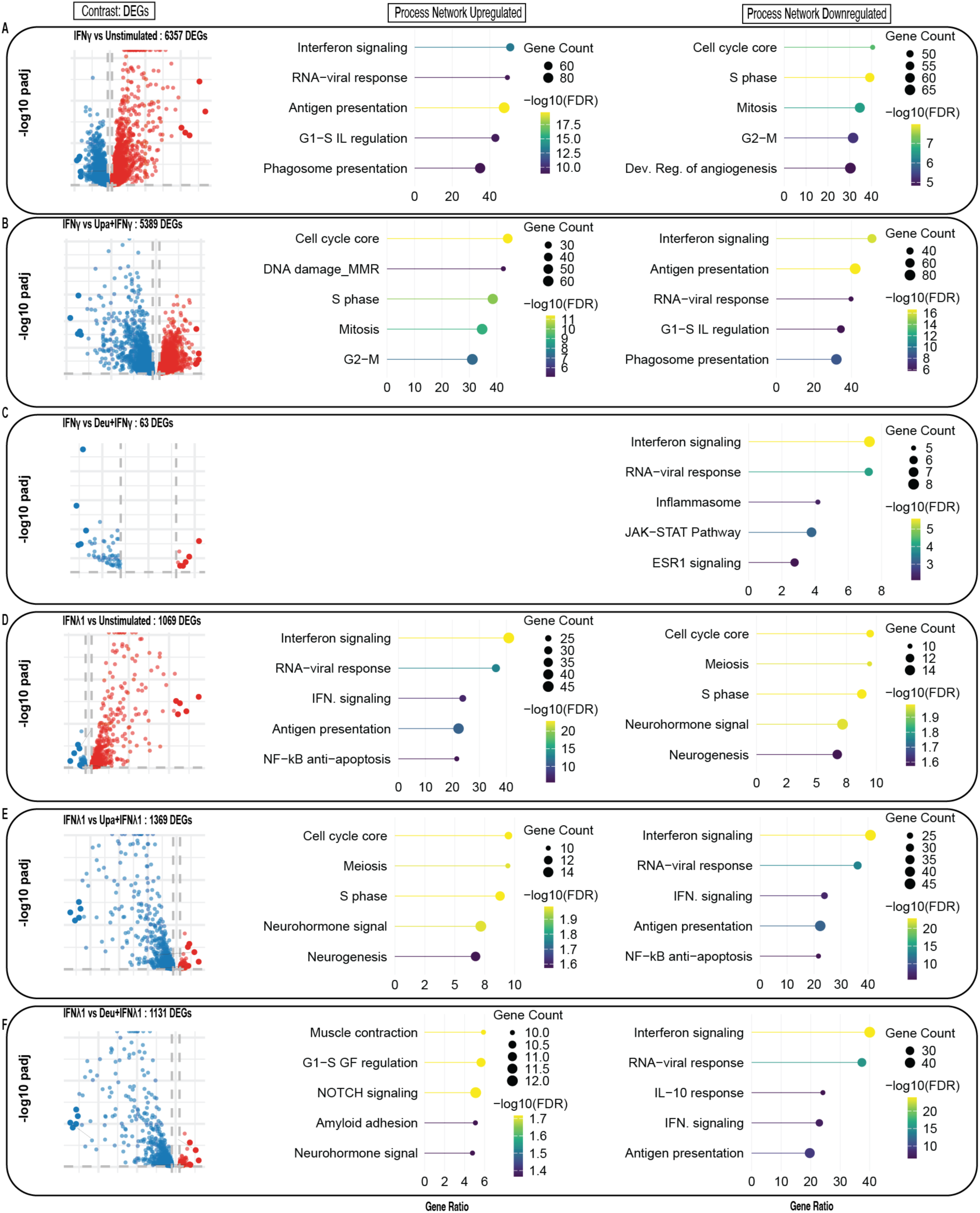
Process enrichment analysis in IFNγ/IFNλ1-stimulated IEOs with or without pretreatment with upadacitinib or deucravacitinib. Volcano plots illustrate differentially expressed genes (DEGs) and corresponding top 5 process networks enriched are shown as lollipop plots for the following comparisons: A) IFNγ vs Unstimulated, B) IFNγ vs Upadacitinib (Upa) + IFNγ, C) IFNγ vs deucravacitinib (Deu) + IFNγ, D) IFNλ1 vs Unstimulated, E) IFNλ1 vs upadacitinib + IFNγ, and F) IFNλ1 vs deucravacitinib + IFNγ. In the lollipop plots, the line length indicates gene ratio (number of DEGs in pathway/total genes in pathway), dot size represents gene counts per process network and color gradient corresponds to statistical significance (-log10(false discovery rate (FDR))). See Supplementary File 5 for processes enriched in response to conditions.

**Supplementary Figure 6:**
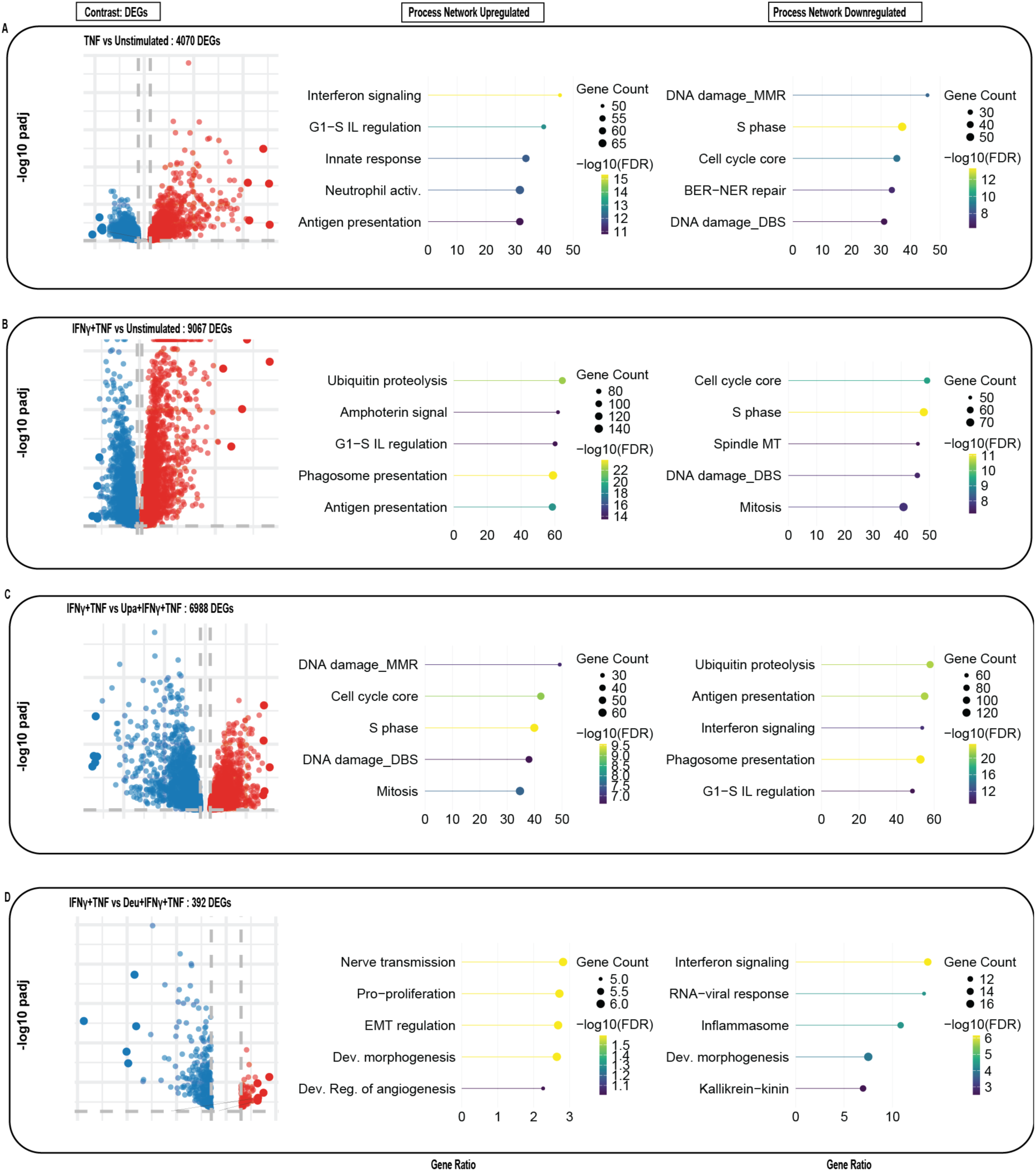
Process enrichment analysis in TNF/IFNγ+TNF-stimulated IEOs with or without pretreatment with upadacitinib or deucravacitinib. Volcano plots illustrate differentially expressed genes (DEGs) and corresponding process networks enrichment analyses shown as lollipop plots for the following comparisons: A) TNF vs Unstimulated, B) IFNγ + TNF vs Unstimulated, C) IFNγ + TNF vs upadacitinib (Upa) + IFNγ + TNF, D) IFNγ + TNF vs deucravacitinib (Deu) + IFNγ + TNF. In the lollipop plots, the line length indicates gene ratio (number of DEGs in pathway/total genes in pathway), dot size represents gene counts per process network and color gradient corresponds to statistical significance (-log10(false discovery rate (FDR))). See Supplementary File 5 for processes enriched in response to different conditions.

**Supplementary Figure 7.**
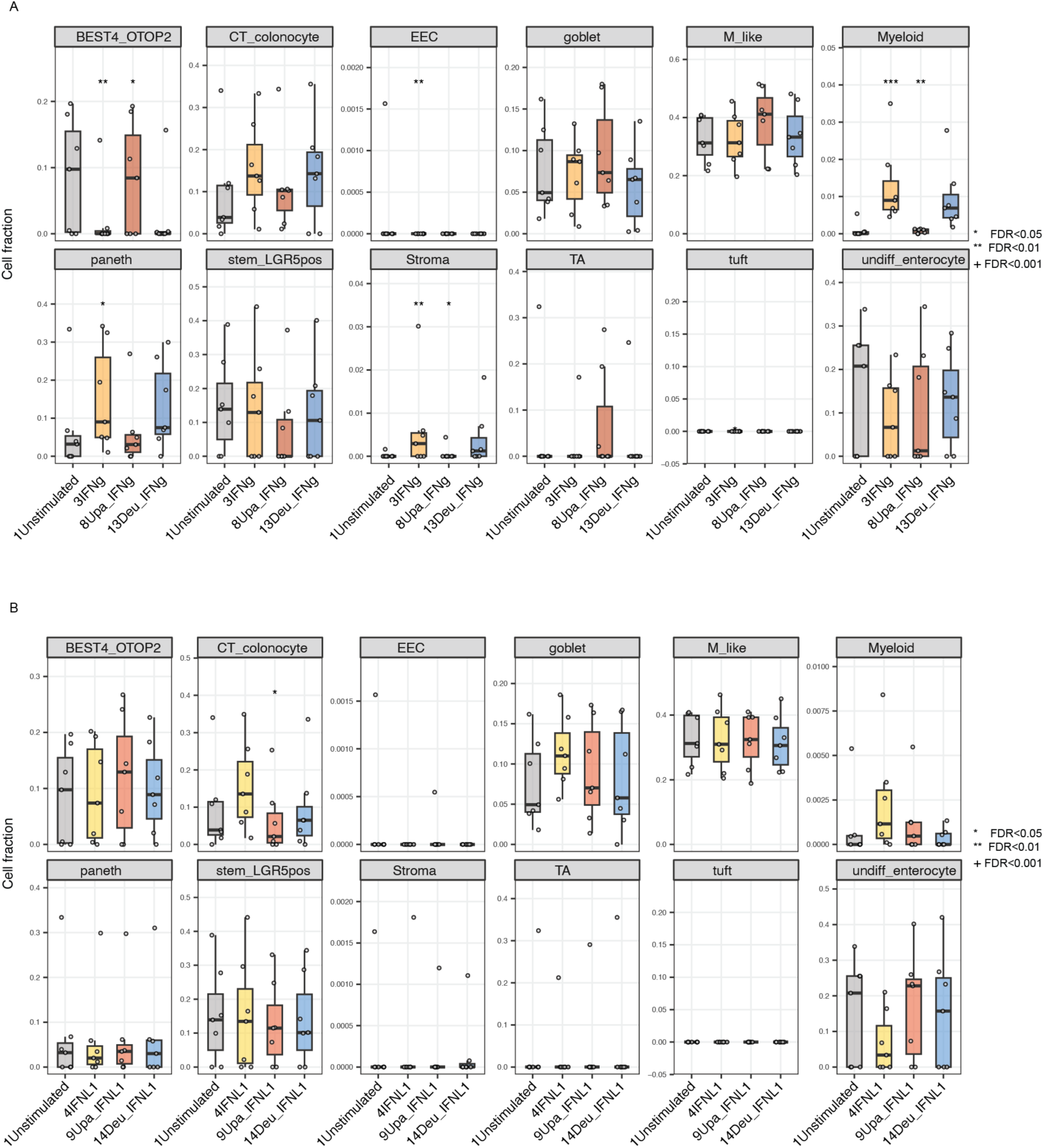
Estimated fractions of epithelial cell types in IEOs under A) IFNγ and B) IFNλ1 stimulation. Box plots show the fractions of differentiated and undifferentiated epithelial subsets (colonocytes, TA, BEST4+/OTOP2+, goblet, paneth, LGR5+ stem, tuft, EEC, M-like, and undifferentiated cells) as well as myeloid and stromal compartments, from 7 donors across the indicated conditions. Significance was assessed by linear mixed models (LMM) on centered log-ratio-transformed proportions with donor as a random intercept, and P-values were adjusted using the Benjamini-Hochberg method. *FDR < 0.05, **FDR < 0.01, ⁺FDR < 0.001. Raw fractions are provided in Supplementary File 6.

**Supplementary Figure 8.**
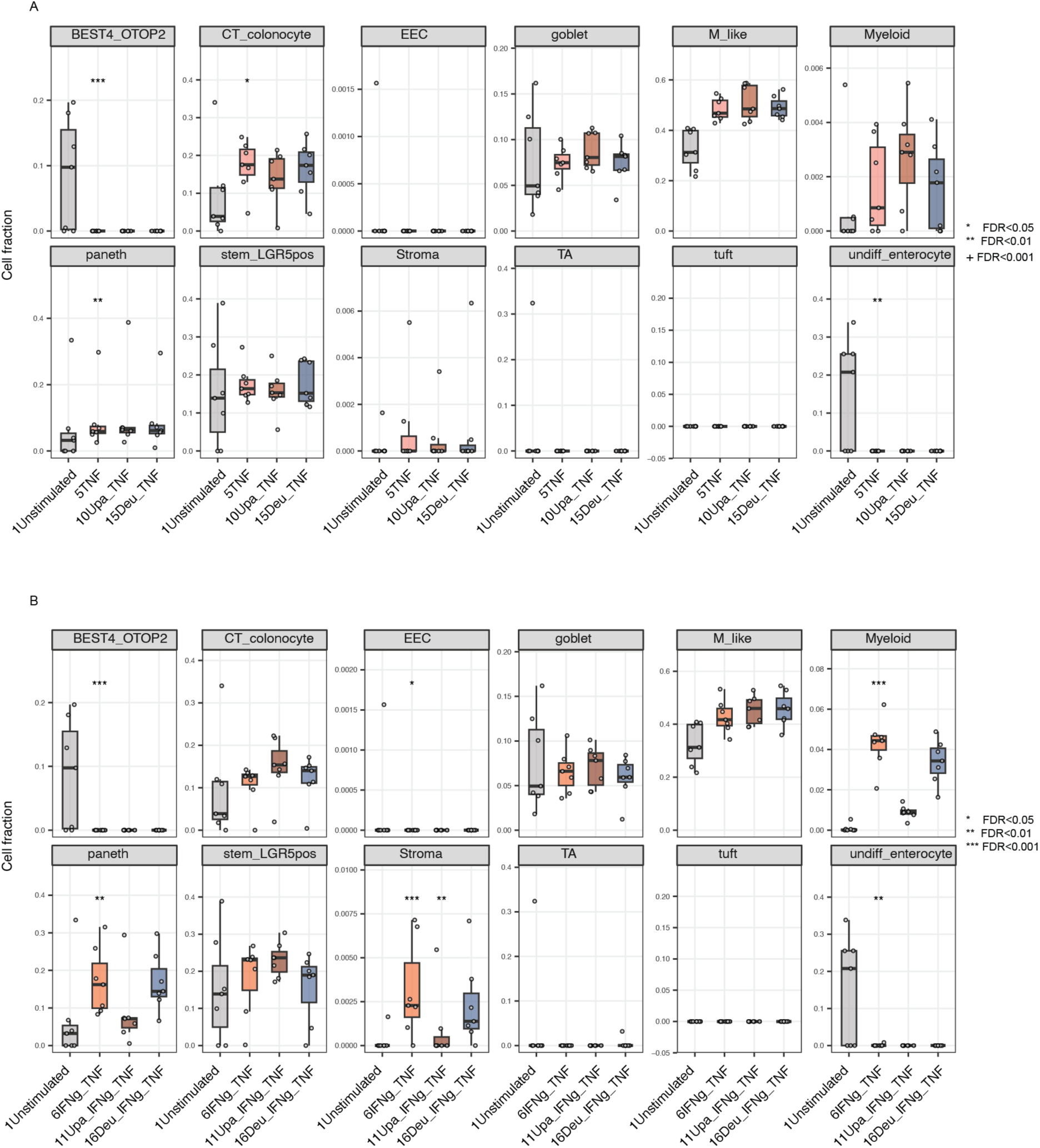
Estimated fractions of epithelial cell types in IEOs under A) TNF and B) IFNγ + TNF stimulation. Box plots show the fractions of differentiated and undifferentiated epithelial subsets (colonocytes, TA, BEST4+/OTOP2+, goblet, paneth, LGR5+ stem, tuft, EEC, M-like, and undifferentiated cells) as well as myeloid and stromal compartments, from 7 donors across the indicated conditions. Significance was assessed by linear mixed models (LMM) on centered log-ratio-transformed proportions with donor as a random intercept, and P-values were adjusted using the Benjamini-Hochberg method. *FDR < 0.05, **FDR < 0.01, ⁺FDR < 0.001. Raw fractions are provided in Supplementary File 6.

**Supplementary Figure 9.**
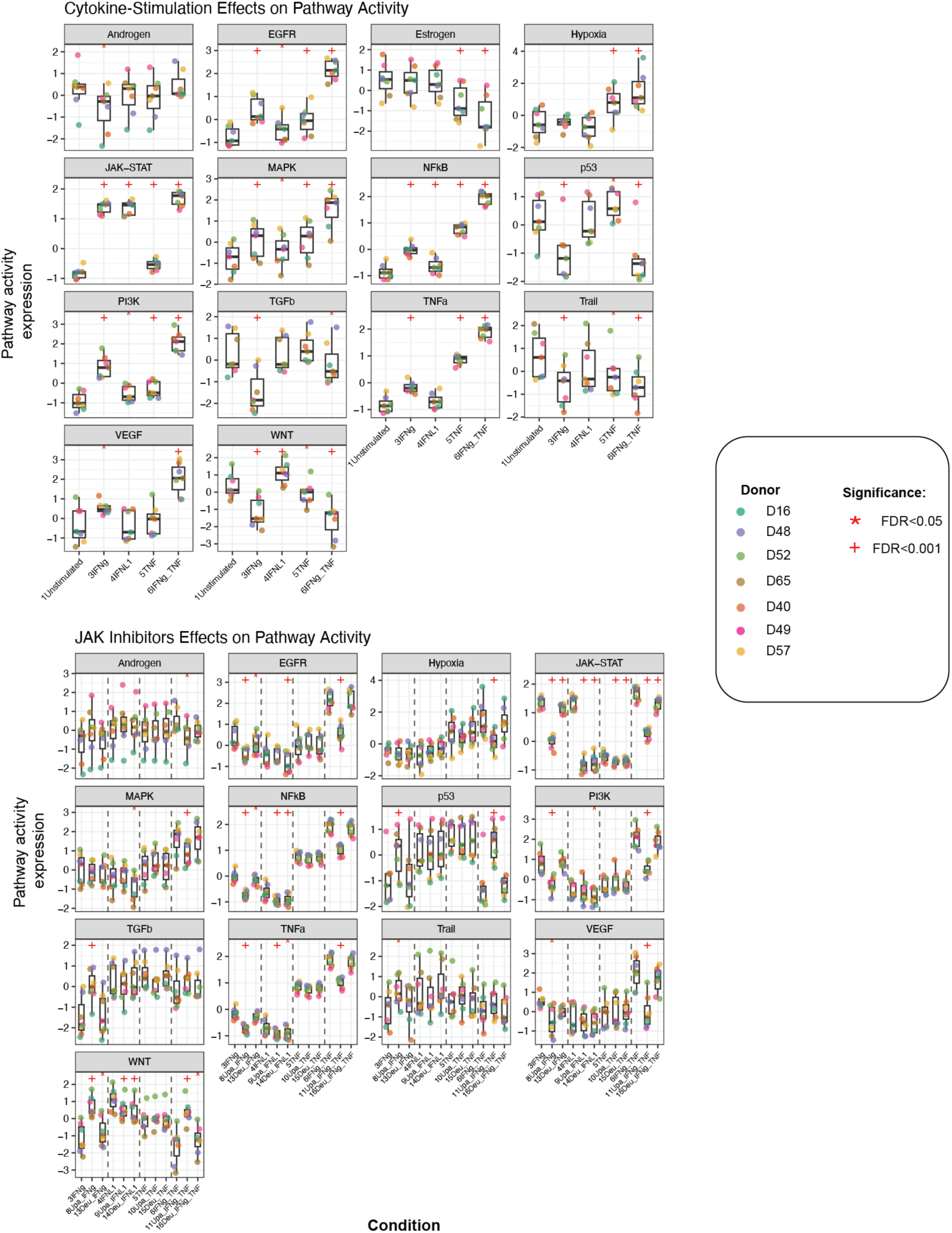
Inferred activity of different pathway signatures in IEOs following JAK/TYK2 inhibitor with or without cytokine-stimulation. Pathway activity was inferred using PROGENy (65), based on variance-stabilized transformed RNA-seq expression data. Box plots display individual pathway activity expression in IEOs (*n* = 7) in A) stimulated conditions compared to unstimulated controls and B) JAK/TYK2 inhibitor pretreatment followed by stimulated conditions compared to their respective stimulated controls as indicated. Statistical significances were assessed using linear mixed model (LMM) with donor as random effect, and P-values were adjusted using Benjamini-Hochberg method for multiple comparisons testing. Upa - Upadacitinib, Deu - Deucravacitinib, *P<0.05, **^+^**P<0.001.

**Supplementary Figure 10.**
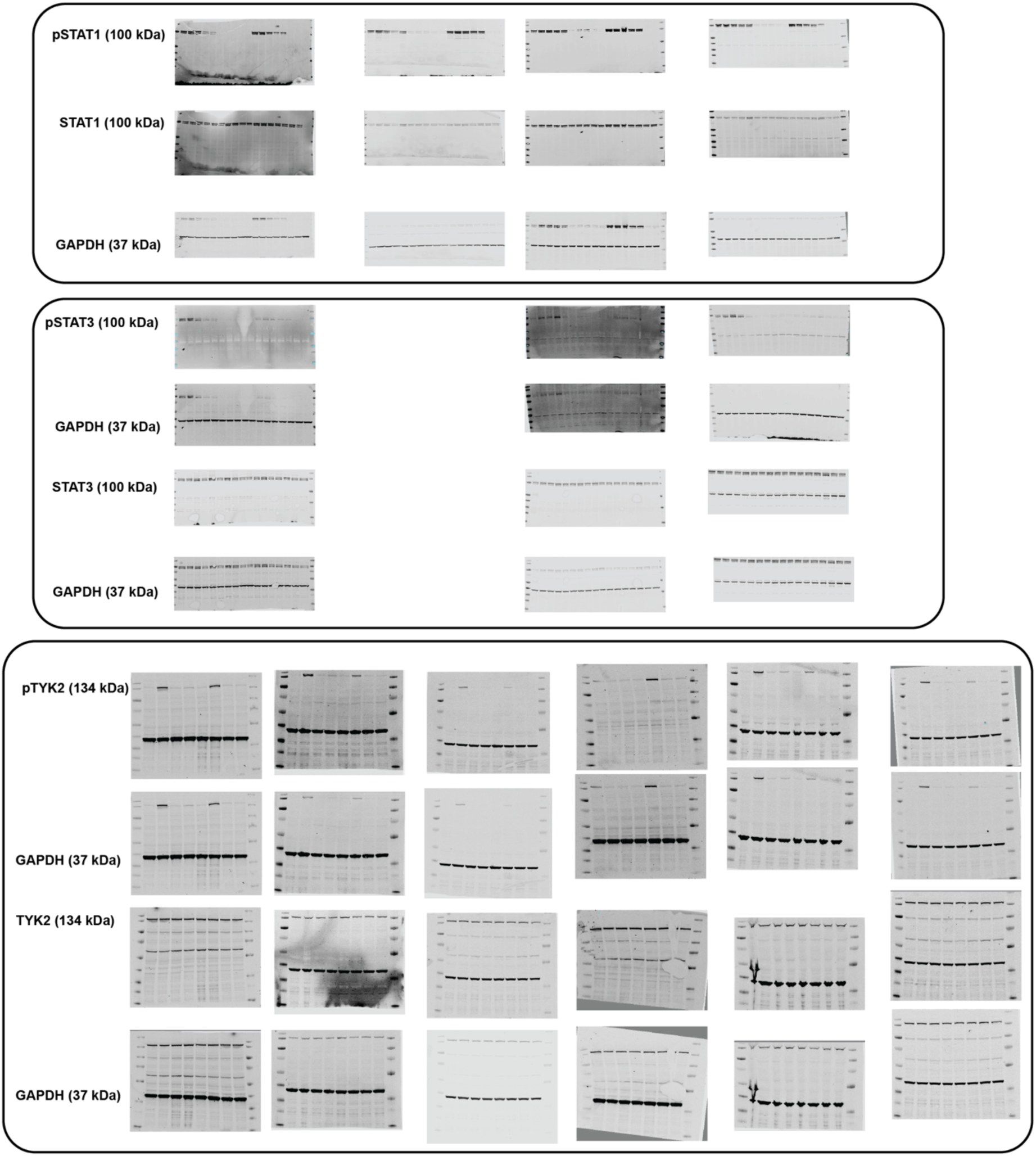
Full blot images from all the replicates of HT-29 cells showing phosphorylation of STAT1, STAT3, and TYK2, corresponding to data shown in Supplementary Figure 1.

**Supplementary Figure 11.**
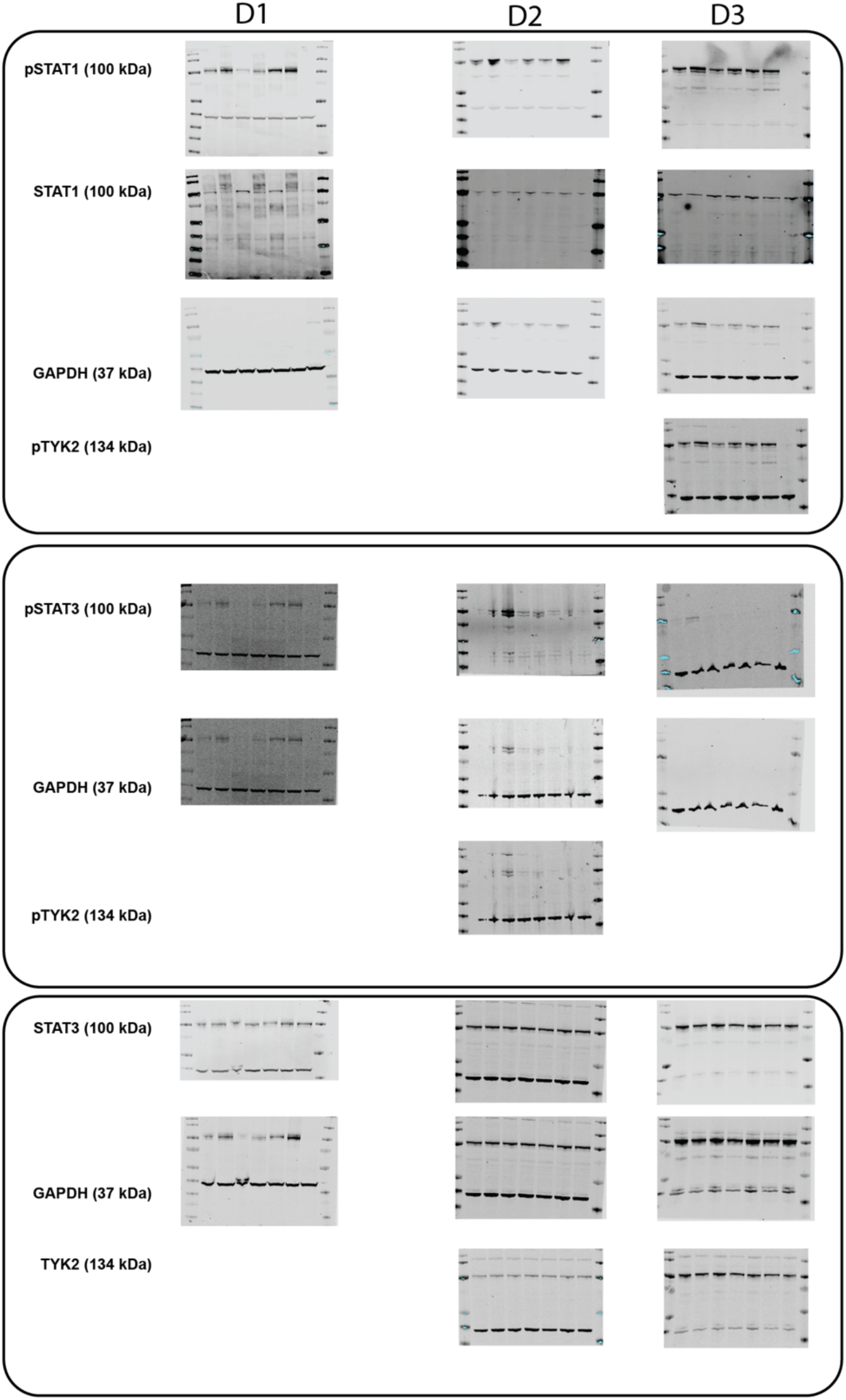
Full blot images from all the replicates of intestinal epithelial organoids (IEOs) stimulated with IFNβ, IFNγ, and IFNλ1, showing phosphorylation of STAT1, STAT3, and TYK2 corresponding to Supplementary Figure 1.

**Supplementary Figure 12.**
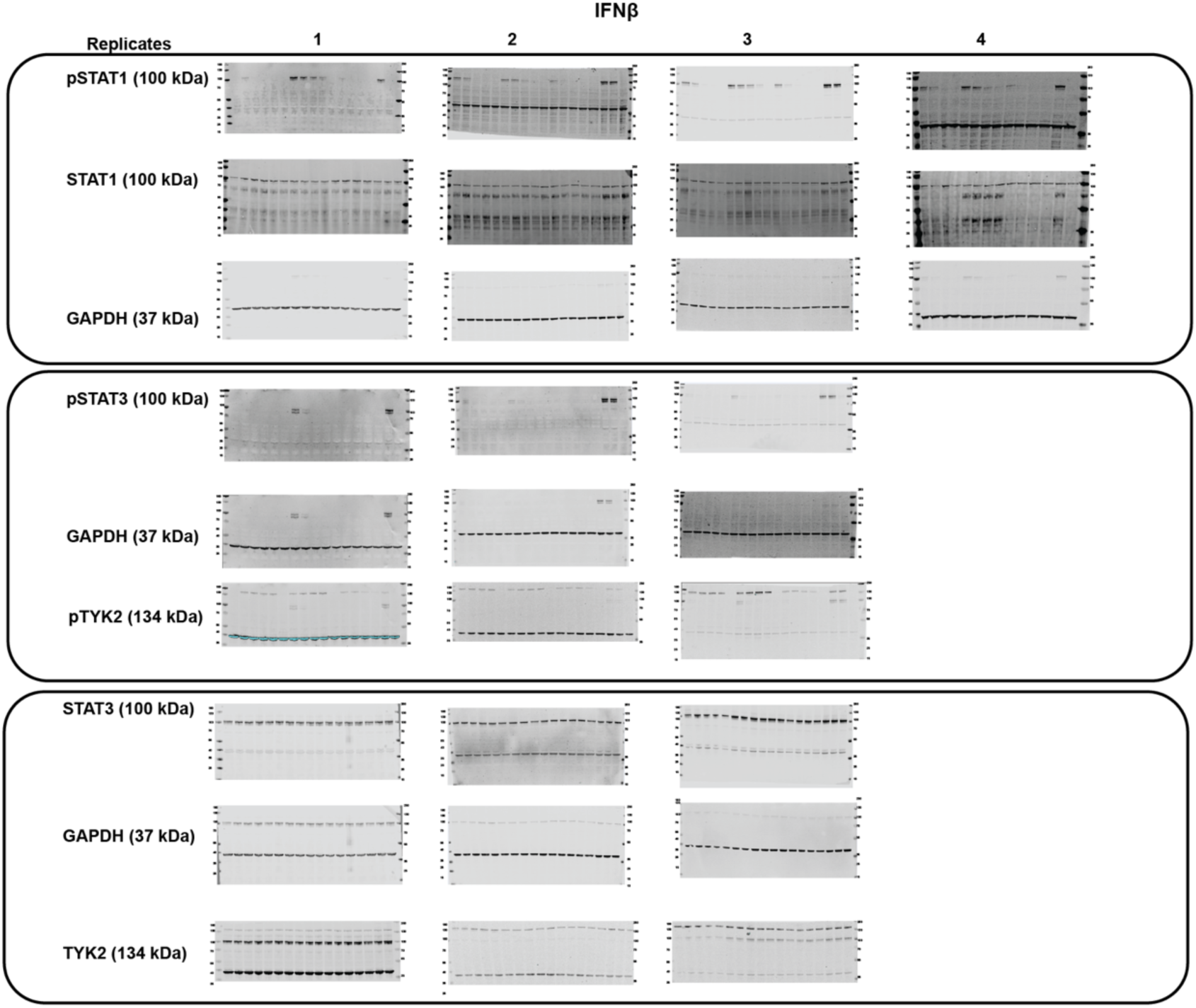
Full blot images from IEOs stimulated with IFNβ following pretreatment with tofacitinib, upadacitinib or filgotinib, showing phosphorylation of STAT1, STAT3, and TYK2 corresponding to Figure 1B and Supplementary Figure 2.

**Supplementary Figure 13.**
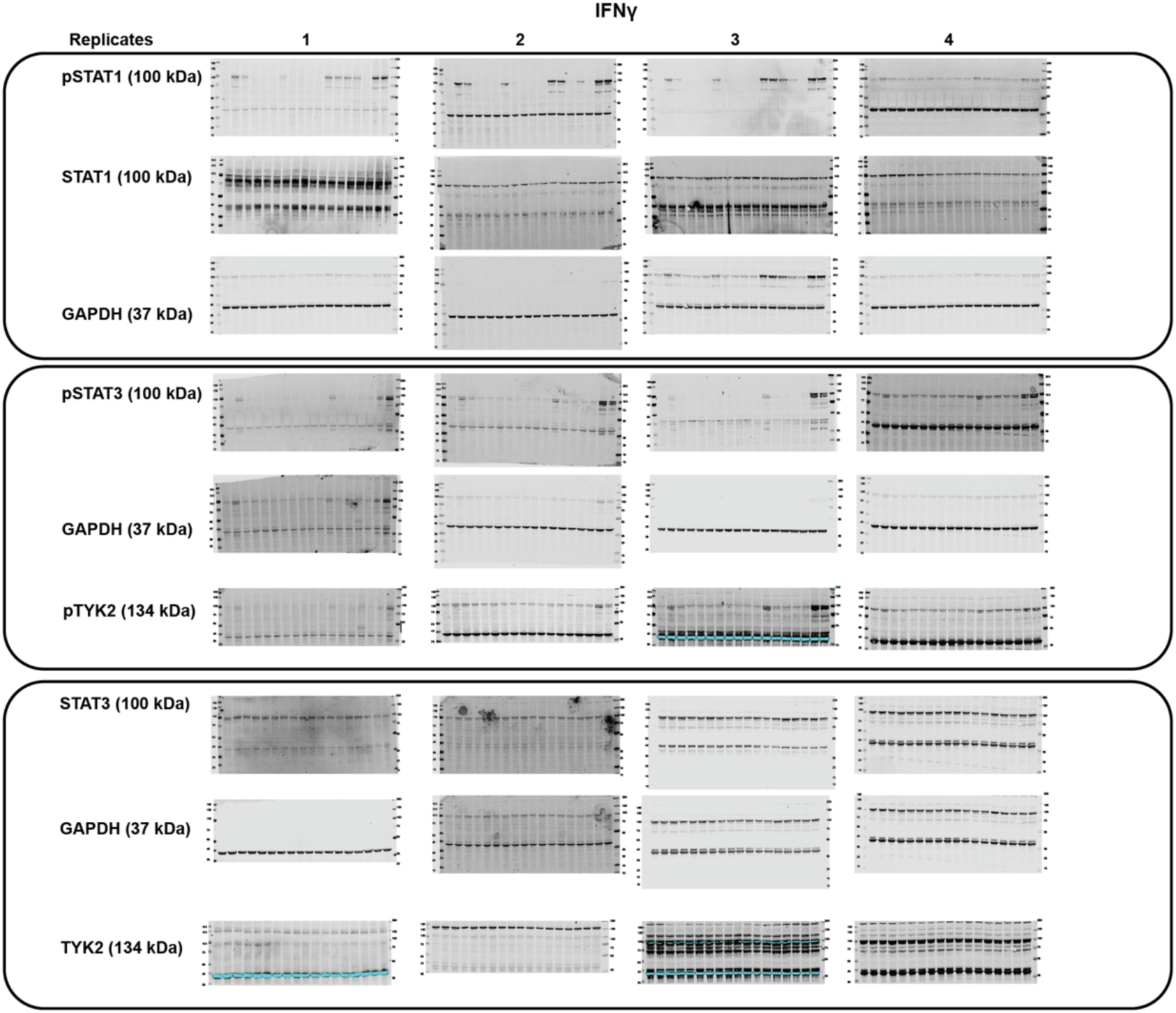
Full blot images from IEOs stimulated with IFNγ following pretreatment with tofacitinib, upadacitinib or filgotinib, showing phosphorylation of STAT1, STAT3, and TYK2 corresponding to Figure 1C and Supplementary Figure 3.

**Supplementary Figure 14.**
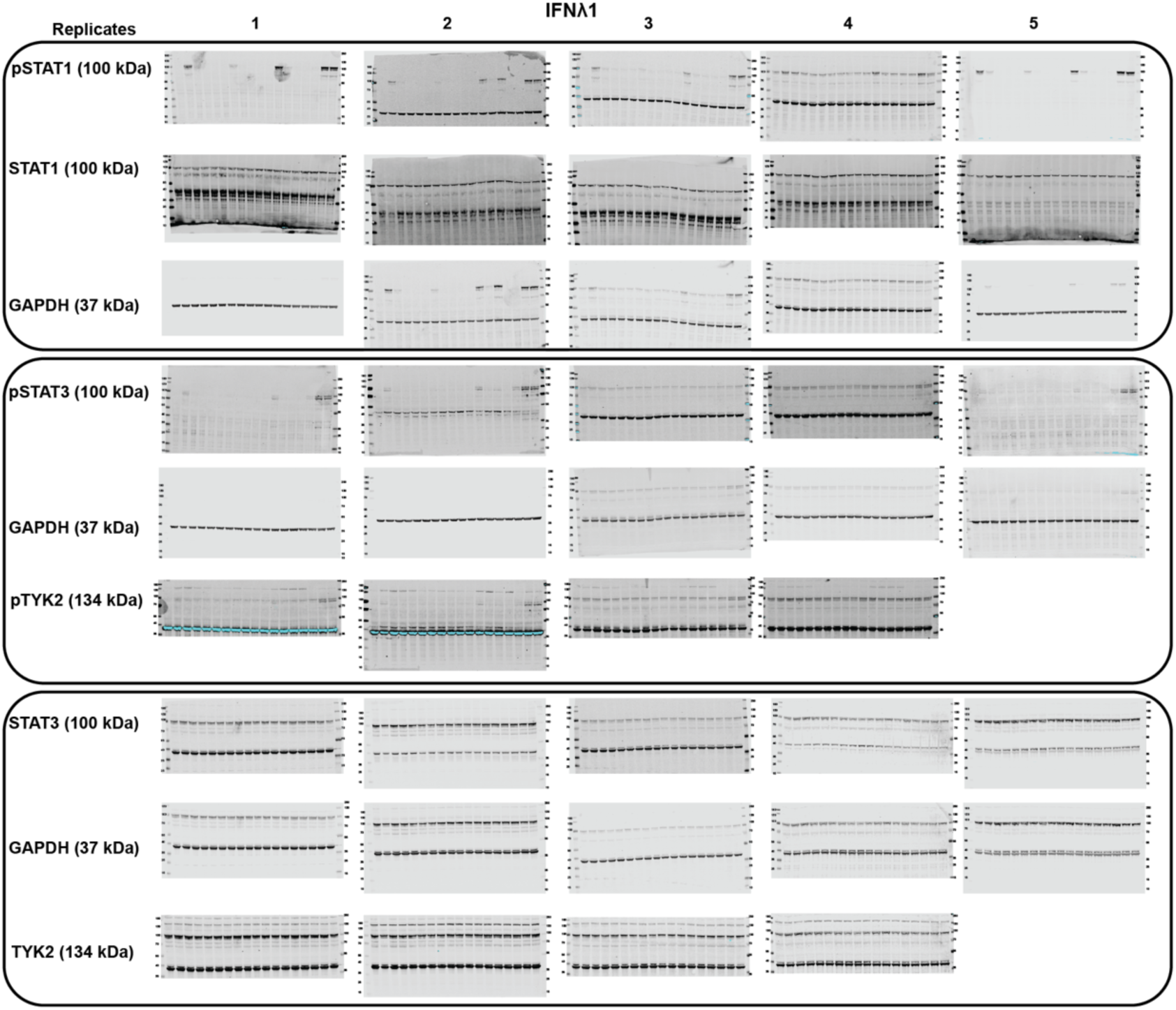
Full blot images from IEOs stimulated with IFNλ1 following pretreatment with tofacitinib, upadacitinib or filgotinib, showing phosphorylation of STAT1, STAT3, and TYK2 corresponding to Figure 1D and Supplementary Figure 4.

**Supplementary Figure 15.**
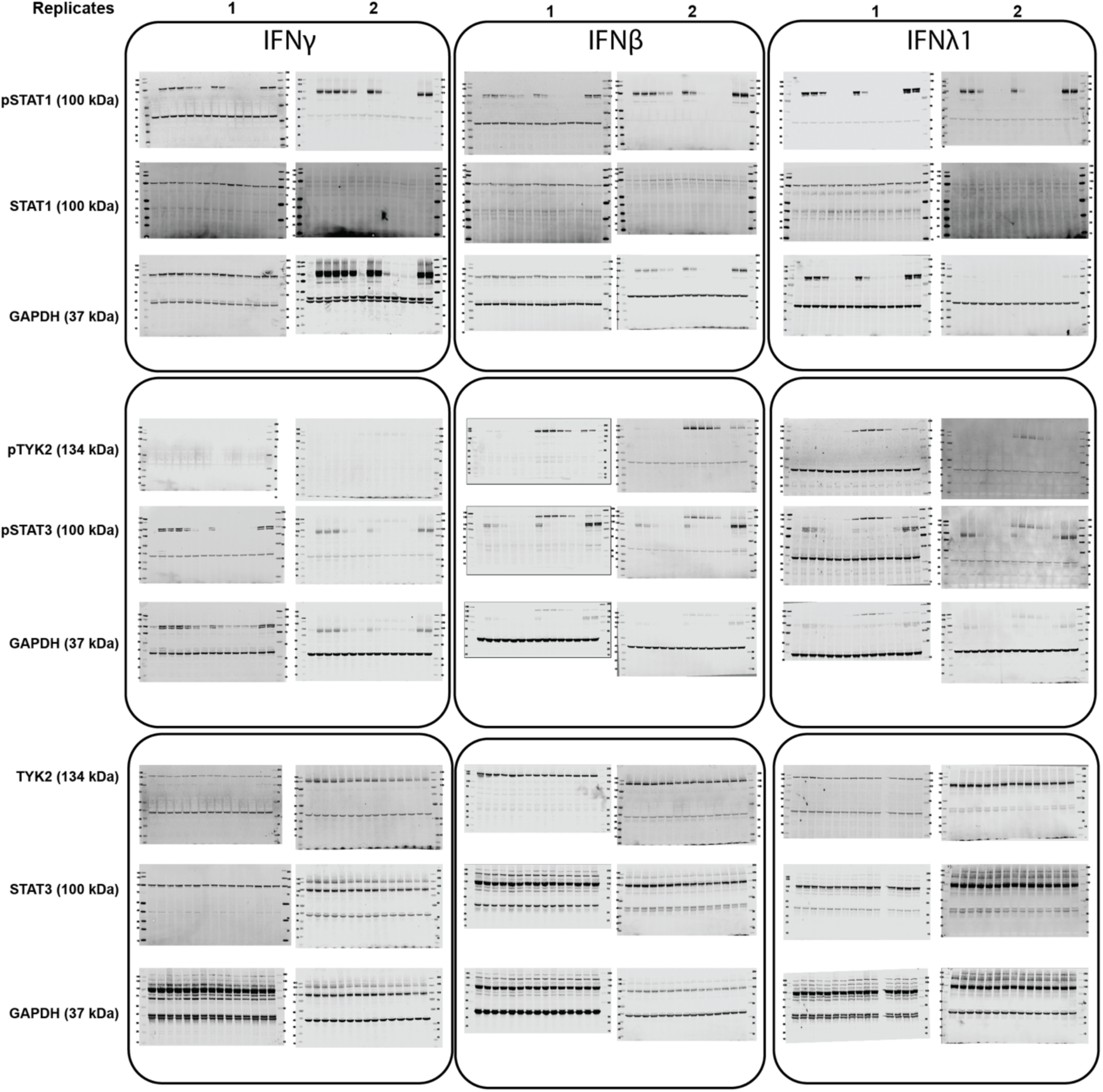
Full blot images from IEOs stimulated with IFNβ, IFNγ, and IFNλ1 following pretreatment with deucravacitinib or brepocitinib, showing phosphorylation of STAT1, STAT3, and TYK2 corresponding to Figure 1B-D and Supplementary Figure 2-4.

